# Sensitive detection of chloroplast movements through changes in leaf cross-polarized reflectance

**DOI:** 10.1101/2023.10.24.563792

**Authors:** Paweł Hermanowicz, Aleksandra Giza, Katarzyna Sowa, Paweł Korecki, Justyna Łabuz

## Abstract

We present a sensitive method for non-contact detection of chloroplast movements in leaves and other photosynthetic tissues, based on changes in the magnitude of cross-polarized reflectance. We examined changes in bidirectional red light reflectance during irradiation with blue light, known to trigger chloroplast relocations. Experiments on the model plant *Arabidopsis thaliana*, wild-type, and several mutants with disrupted chloroplast movements showed that the chloroplast avoidance response, induced by high blue light, led to a substantial increase in diffuse reflectance of unpolarized red light. The effects of the accumulation response in low blue light were the opposite. The specular reflectance of the leaf was unaffected by the chloroplast positioning. To further improve the specificity of the detection, we examined the effects of chloroplast relocations on the leaf reflectance of a linearly polarized incident beam. The greatest relative change associated with chloroplast movements was observed when the planes of polarization of the incident and detected beams were perpendicular. Further experiments revealed that the chloroplast positioning affected the magnitude of depolarization of light by the leaf. We applied the developed approach to examine chloroplast relocations in four angiosperm species collected in the field. The method allowed us to detect the chloroplast avoidance response in the green stems of bilberry, a sample not amenable to transmittance-based detection. Despite the importance of chloroplast movements for the optimization of photosynthetic efficiency and biomass production, high throughput reflectance-based methods are not routinely used for their detection. This method opens the possibility of non-invasive, non-contact detection of chloroplast relocations in a manner insensitive to the orientation of the leaf.

## 1. Introduction

Chloroplast movements allow plants to achieve high photosynthetic performance and biomass production in dynamic light conditions (Hart et al., 2019). In low light, chloroplasts gather at cell walls perpendicular to the light direction to maximize photosynthetic efficiency. In high light, chloroplasts move to walls parallel to the direction of light as a photoprotective mechanism. In flowering plants, chloroplast relocations are triggered by blue light, perceived through UV/blue light receptors phototropins (Banaś et al., 2012), (Łabuz et al., 2022). Two phototropins control chloroplast movements in the model plant *Arabidopsis thaliana*. Phot1 and phot2 trigger chloroplast accumulation (Sakai et al., 2001), while sustained chloroplast avoidance is elicited only by phot2 (Jarillo et al., 2001), (Kagawa et al., 2001).

Initially, chloroplast relocations were examined using light microscopy, as in the pioneering work of (Senn, 1908). Using modern microscopic techniques, tracking of movement of a single chloroplast within a cell in response to microbeam irradiation is possible (Kagawa & Wada, 2000), (Tsuboi & Wada, 2010), (Wada, 2013). Fixed leaf cross-sectioning allows for studying chloroplast positioning in multiple mesophyll layers but precludes time course studies (Higa & Wada, 2016). Chloroplast movements cause substantial changes in the optical properties of the leaf, affecting its transmittance and reflectance (Zurzycki, 1961). As red light usually does not induce the movements in angiosperms, it is used to monitor changes in leaf optical properties triggered by actinic blue irradiation. The most widely used approach to detect chloroplast relocations consists of the recording of light-induced transmittance changes, both in the laboratory (Inoue & Shibata, 1974) and in field conditions (Williams et al., 2003). The transmittance-based detection is robust and repeatable provided that an integrating sphere or the opal method (Shibata, 1958), (Walczak & Gabrys, 1980) is employed to eliminate the dependence of the signal on the sample-detector distance. Transmittance-based methods require direct access to the leaf, which limits their throughput and applicability in field conditions. Approaches using a multi-plate reader have been employed to overcome some of the limitations and enable rapid analysis of many samples (Johanssson & Zeidler, 2016). An alternative method to detect chloroplast relocations is through measurements of the changes in leaf reflectance. An increase in leaf reflectance accompanying the avoidance response has been observed in *Begonia multiflora* (Seybold, 1955) and *Lemna trisulca* (Zurzycki, 1961). The detection of chloroplast movements through changes in light reflectance has been reported by (Park et al., 1996), (Davis et al., 2011), (Baránková et al., 2016), (Dutta et al., 2015), (Dutta et al., 2017). Red light leaf reflectance has been used to monitor chloroplast relocations in *Arabidopsis thaliana* during light treatments in a growth chamber (Dutta et al., 2015). Measurements of diffuse white light leaf transmittance and reflectance, performed for several species using an integrating sphere, revealed that cell shape substantially affects the ability of chloroplasts to move, influencing blue-light-induced changes in leaf absorptance and scattering (Davis et al., 2011). A detailed study of blue-light-induced changes in leaf transmittance and reflectance, using an integrated sphere, was also performed in *Nicotiana tabacum* (Baránková et al., 2016). Interestingly, it revealed that changes due to chloroplast movements in leaf absorptance of a collimated measuring beam were substantially greater when the beam illuminated the adaxial as opposed to the abaxial side. The differences disappeared when the leaf was infiltrated or the beam was diffused, pointing to the significance of refraction at the cell–airspace interface. In our previous work (Hermanowicz & Łabuz, 2024), we showed that the increase in the leaf reflectance due to the chloroplast avoidance response in *Arabidopsis* leaves could be observed in the whole visible range, especially in the 580 - 660 nm region. Spectra extracted from the hyperspectral images of leaves from lab-grown *A. thaliana* and *Nicotiana benthamiana*, recorded at small incidence and viewing angles, could be accurately classified according to their chloroplast positioning using a machine learning methods, though good results were also obtained with a simple vegetation index. Effects of chloroplast positions on the reflectance spectrum might bias estimates of other plant traits based on spectral imagery and vegetation indices (Hermanowicz & Łabuz, 2024).

An important drawback of reflectance-based measurements of plant traits is the dependence of the amount of reflected light reaching the detector on the orientation of the leaf (Li et al., 2018). Specular reflectance exhibits strong angular dependence, hindering chloroplast movement detection, as reported in *Alocasia macrorrhiza* (Gorton et al., 1999). Several potential approaches can be proposed to alleviate the effect of the leaf orientation on the reflectance-based chloroplast movement detection, which may be used in concert: 1) simultaneous detection of reflectance at several angles of observation, 2) measuring the plant or canopy geometry in parallel to reflectance measurements, 3) incorporation of polarization information. This study aimed to determine whether the specificity of chloroplast movement detection could be improved by analyzing the polarization of light reflected from the leaf. Unpolarized light specularly reflected by the wax layer or the cuticle becomes partially polarized, with the degree and the plane of polarization, dependent on the orientation of the leaf. By contrast, light diffusely reflected from the leaf interior, through scattering and multiple refractions at the border between intercellular spaces and mesophyll cell walls, does not undergo substantial polarization (Vanderbilt et al., 1985). This suggests that by separating the polarized and non-polarized components of the reflected light, the detection of chloroplast movements independently of the spatial orientation of the leaf would be feasible. The diffuse and specular reflection mechanisms also differ in their ability to depolarize light. When the leaf is illuminated with a polarized beam, light reflected from the leaf surface does not undergo substantial depolarization. By contrast, diffuse reflectance from the leaf interior undergoes considerable depolarization (Rvachev & Guminetskii, 1966), (Brakke, 1994), likely due to multiple scattering. This suggests that the cross-polarized reflectance may contain information about the state of the leaf mesophyll, particularly the positioning of chloroplasts. Previously, the magnitude of depolarization was used to remotely sense the nitrogen status of crops (Kalshoven et al., 1995), possibly due to the nitrogen-availability effects on the chlorophyll amount.

To examine the link between chloroplast positioning and polarized red light reflectance, we used several lines of the model plant *Arabidopsis thaliana*: wild-type Columbia and photoreceptor mutants, *phot1, phot2* (lacking sustained chloroplast avoidance), and *phot1phot2* (disrupted chloroplast movements). We also employed a signaling mutant, *jac1,* deficient in chloroplast accumulation (Suetsugu et al., 2005). These mutants were in the Columbia background, characterized by leaves covered with trichomes. To investigate whether the presence of trichomes affects the performance of the reflectance-based detection of chloroplast relocations, we also examined the *glabra1* mutant. To evaluate the extent of applicability of the developed methods, we assessed their performance in photosynthetic organs of several wild species.

## 2. Materials and Methods

### 2.1. Plant material

WT Col-0 *Arabidopsis thaliana* (L.) Heynh., *glabra1, phot1* (SALK_088841) (Lehmann et al., 2011), *jac1-3* (WiscDsLox457-460P9) (Hermanowicz et al., 2019), *phot2* (Jarillo et al., 2001), *phot1phot2* (Hermanowicz et al., 2019), were sown in Jiffy-7 pots (Jiffy Products International AS) and stratified at 4°C for 2 days. Plants were cultivated in a growth chamber (Sanyo MLR 350H) at 23°C, 80% relative humidity, a photoperiod of 10 h light and 14 h darkness, at photon irradiance of 70 (±15) μmol·m^−2^·s^−1^ (measured at the plane parallel to the rosettes) supplied by fluorescent lamps. The position of plants from different lines in the chamber was randomized and the trays with plants were rotated 6 – 7 times per week. The experiments were performed on rosette leaves detached from dark-adapted 5-week-old plants. Leaf chlorophyll content was assessed with the Leaf State Analyzer (Walz, Germany).

Wild plants were collected in March of 2023 and 2024. Plants were harvested in the afternoon, kept dark-adapted in water, and measured the next day. Flowering specimens of *Draba verna* L. (common whitlowgrass) were collected in a xeric meadow covering the south-facing slope of Górka Pychowicka hill (Kraków, 50°1’45" N 19°53’6" E). Chloroplast movements were induced by irradiation of the adaxial side of rosette leaves (Fig. 7 A)*. Stellaria media* (L.) Vill. (chickweed) was collected in a young birch-alder forest near Jagiellonian University Campus in Kraków, Poland (50°1′44″ N 19°54’6" E). The adaxial surface of the third leaf from the top of the plant (asterisk in Fig. 7 B) was irradiated. *Tussilago farfara* L. (coltsfoot) was collected at human-disturbed sites (lawns and construction sites) near the Jagiellonian University Campus. Chloroplast movements were induced by irradiation of the abaxial (sun-exposed) side of the scales covering flowering shoots (Fig. 8 A). *Vaccinium myrtillus* L. (bilberry) stems were collected in a fir forest on the north-facing slope of Mnisko Mt in the Gorce Range, Poland (750 - 800 m a.s.l, 49°34’40” N 20°2’39" E). Chloroplast movements were examined in terminal branches (asterisk in Fig. 8 B).

### 2.2. Definitions and conventions

We will use the term bidirectional reflectance *R*(θ*_i_*,φ*_i_,*θ*_v_*,φ_v_,) to refer to the ratio between the radiance of the sample when viewed along the direction (θ*_v_*,φ_v_), and irradiance received from the direction (θ*_i_*,φ*_i_*) (Nicodemus, 1965), (Schaepman-Strub et al., 2006), where θ and φ are polar and azimuthal angles, respectively. As in (Vanderbilt et al., 1985), we calculate polarized reflectance as the ratio of the polarized reflectance and part of the irradiance corresponding to the appropriate polarization state, even when the incident radiation was unpolarized. In experiments with polarized incident light, total reflectance Δ*R_tot_* was calculated as the sum of reflectance values obtained for two orthogonal orientations of the analyzer (the polarizer in front of the detector). Light polarized in the plane of the normal and the incident ray is referred to as *P*-polarized. Light polarized in the plane perpendicular to the plane of incidence is described as *S*-polarized. In our experiments, the extrema of detected signal intensity for light reflected from a leaf coincided with the *S* and *P* orientations. Thus, the degree of linear polarization (DoLP) of reflected light was estimated as the absolute value of (*R_s_ - R_P_*)/(*R_s_*+ *R*_P_), where *R_s_* and *R_P_* are reflectance values obtained with the *S* and *P* orientations of the polarizer, respectively. Our convention for specifying angles of incidence and observation is shown in Fig. 1 A. As the incident rays, the leaf normal, and the direction of observation were all contained in the same plane, we specify the directions of incidence and observation using a single angle, measured with respect to the leaf normal. Thus, a positive angle α corresponds to the polar angle α and azimuth of 0°, while a negative angle -α describes the same direction as the polar angle α and azimuth of 180°.

**Fig. 1.**
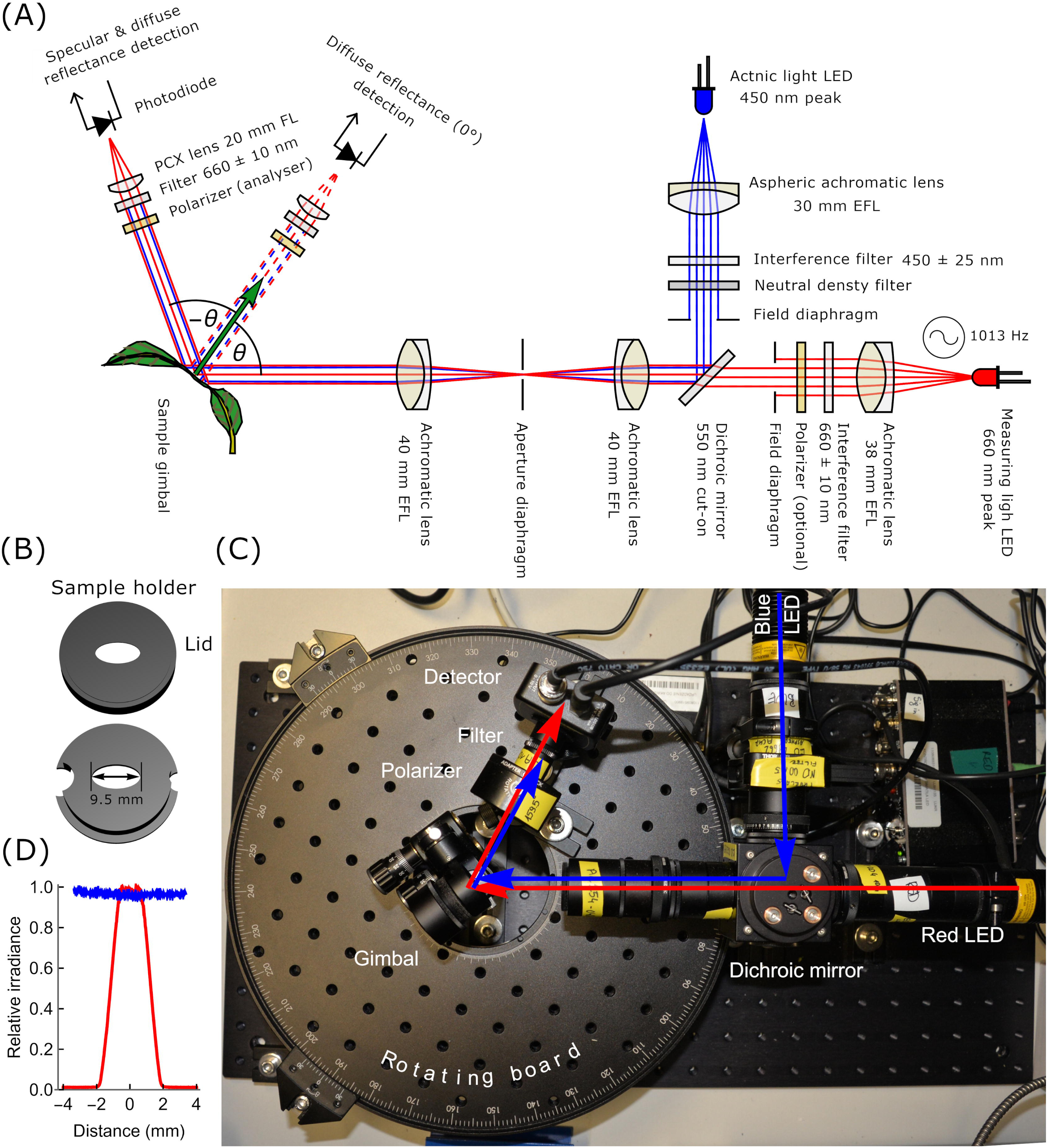
(A) Schematic representation of the reflectometric set-up used in the study. Blue and red beams are collimated, combined using a dichroic mirror, and directed towards the leaf. Red light, modulated at 1013 Hz, is used to monitor reflectance changes due to chloroplast movements. Blue, continuous light induces chloroplast movements. The angle of incidence θ of the combined blue and red beams onto the surface of the leaf is either 40° or 55°. The directional reflectance is measured at a variable angle of observation, with the detector lying in the plane spanned by the direction of incident rays and the normal to the leaf. Reflected light passes through a linear polarizer, an interference filter (blocking blue actinic light and attenuating chlorophyll fluorescence), and a plano-convex lens, which focuses reflected light on the detector. (B) A model of the 3D printed sample holder used to mount the leaf in the gimbal. The holder features a hole to prevent reflection from the material beneath the leaf. (C) Photo of the set-up. The rotating board provides control over the polar angle at which reflected light is detected. (D) The relationship between the blue and red beam relative irradiances on the distance from the beam centre.

### 2.3. Optomechanical setup for measurement of blue light-induced changes in leaf bidirectional reflectance

The measuring beam was produced by a red LED (nominal wavelength of 660 nm, manufacturer part number M660L4, Thorlabs) and collimated with an achromatic lens (#49778, Edmund Optics). The collimated beam was passed through a bandpass interference filter (660 nm max, 10 nm FWHM, #15-126, Edmund Optics) to reduce the width of its spectrum. In some experiments (Fig. 4 - 5), a linear polarizer (40 dB extinction ratio, #47316, Edmund Optics) was inserted next to the filter. The actinic beam was supplied by a blue LED (nominal wavelength of 455 nm, M455L4, Thorlabs), collimated with an aspherized achromatic lens (#49-662, Edmund Optics), and passed through a bandpass interference filter (450 nm, 25 nm FWHM, #15146, Edmund Optics). The spectra of the beams measured at the leaf mount are shown in Fig. S1. The beams were combined with a dichroic mirror (DMLP550, Thorlabs) and passed through a condenser composed of two achromatic lenses (AC 254-040-A, Thorlabs) and an iris diaphragm. The beam uniformity was examined with a CDC camera (Axiocam MRm rev3, Carl Zeiss, Jena). The standard deviation of irradiance of the blue beam within the area probed with red light was equal to ca. 1% of the average irradiance (Fig. 1 D). The size of the beams was controlled with field diaphragms. The radius of the red beam at the sample plane was set to 1.5 mm, to avoid illuminating the midvein, while its power was ca. 29 µW (when unpolarized) or ca. 13 µW (when polarized). The radius of the blue beam was 6 mm. The field of view of the detector (ca. 12 mm in radius) exceeded the size of the red spot. Thus, the detected signal was proportional to radiant intensity and was divided by the cosine of the observation angle to obtain radiance. The leaf was placed in a custom-made holder (Fig. 1 B) mounted in a gimbal (GMB1, Thorlabs), which allowed for control of the angle between the leaf surface and the direction of incident light. The holder consisted of two parts, both with an elliptical hole in the middle, between which the leaf fragment was inserted. Edges of the specimen were covered with wet tissue paper. The angle between the incident beam and the leaf normal was kept at 40° or 55°. The angle of 55° corresponds to Brewster’s angle for the interface between air and a sample made of material with a refractive index of 1.43, close to the values reported for epicuticular waxes (Jacquemoud & Ustin, 2019) (p. 72). When the angle of incidence is equal to Brewster’s angle, the ray reflected specularly off a nonadsorbing, homogenous, and isotropic material (Meeten, 1997) is completely polarized, in the *S* plane, even when the incident beam is unpolarized. In the case of a scattering sample, such as a leaf, light is also reflected diffusely, in all directions, including the direction corresponding to the specular reflection. In our experiments, the reflected light was collected at the angle of observation of 0° (diffuse reflection) and either -40° or -55° (diffuse and specular reflection) with respect to the leaf normal. It passed through a linear polarizer (#47316, Edmund Optics) and a bandpass interference filter (660 nm, #15-126, Edmund Optics) blocking actinic light and most of the chlorophyll fluorescence. The filtered light was focused on a photodetector (amplified silicon photodiode, PDA100A2, Thorlabs) with a plano-convex lens (LA1074-A, Thorlabs). Using an integrating sphere (2P4/M, Thorlabs) to obtain a polarization-insensitive reference detector (McClain et al., 1995) we found the sensitivity of this photodiode to vary by 0.1% with the polarization plane orientation. The angular size of the clear aperture of the collecting lens with respect to the sample center was 0.019 steradian (calculated using our ray-tracing Mathematica package https://github.com/plantPhotobiologyLab/openRayTracer). To control the observation angle, the detector was mounted at the rotation stage (RBB300A/M. Thorlabs). The experiments were performed with a polarizer in front of the red LED either present or removed. When it was removed, the degree of linear polarization of the measuring beam at the sample plane was ca. 0.02. We will refer to this as unpolarized light. The second polarizer, i.e. the analyzer in front of the detector, was always installed. Its transmission axis was either set to parallel (transmits *P*) or perpendicular (transmits *S* component) to the plane of incidence. The LEDs supplying the measuring and the actinic beams were controlled with two programmable current sources (Cyclops drivers, (Newman et al., 2015)) that enabled precise regulation of light power. The measuring beam was amplitude-modulated at 1013 Hz using a sinusoidal signal generated with a field-programmable gate array (FPGA) board (STEMLab 125-14, Red Pitaya, Slovenia). The signal from the detector was fed into the FPGA board, running a lock-in amplifier program (Luda et al., 2019). Modulation of the amplitude of the measuring beam, in combination with the lock-in amplification (detection of a single frequency), improves the signal-to-noise ratio. In our setup, the additional benefit of lock-in amplification was a further reduction of the risk of detection of reflected unmodulated blue light. The actinic beam was controlled from a PC running BeamJ, our publicly available Java application (https://github.com/pawelHerm/beamJ). The application was designed to control custom optomechanical systems for the analysis of light-induced changes in the optical properties of the leaves. A silver mirror (#49194, Edmund Optics) placed in the gimbal was used to record the detected signal produced when the radiant flux incident on the sample was completely reflected. A white, diffusely reflecting polytetrafluoroethylene (PTFE) (Weidner & Hsia, 1981) disk (SM05CP2C, Thorlabs) was also measured whenever the setup was calibrated. However, the reported values of bidirectional reflectance are absolute (not divided by the PTFE signal). Axes of polarizers were determined using the two-polarizer method of (Rowell et al., 1969). The axis of the analyzer was additionally checked using light that underwent *S*-polarization upon reflection from a glass window (#83375, Edmund Optics), placed in the sample gimbal, and illuminated with the measuring beam at Brewster’s angle. Leaf reflectance curves were processed with a custom-written Mathematica (Wolfram Research, USA) script.

### 2.4. Measurements of blue-light-induced changes in leaf hemispherical transmittance

Hemispherical leaf transmittance changes due to chloroplast movements were assessed with the photometric method (Walczak & Gabrys, 1980), (Gabryś et al., 2017), using a custom-built photometric setup. The device used the same blue and red LED sources as in the reflectometric setup. The detached leaf was mounted in front of a port of an integrating sphere (IS200-4, Thorlabs). The signal was detected with a photodiode detector (DET100A2, Thorlabs), which was mounted at another port of the sphere. Light reaching the detector was filtered with an absorptive/interference compound filter (660 nm max, BN660, Midwest Optical Systems, USA) to reduce shifts in the transmission band of an interference filter.

### 2.5. Light measurements

Chloroplast movements were induced with either 0.1 µmol m^−2^ s^-1^ (accumulation) or 100 µmol m^−2^ s^-1^ (avoidance) of blue light. The spectra of actinic and measuring light used in the reflectometric setup were measured with a BLACK Comet SR spectrometer (25 µm slit, Stellar Net, USA), equipped with a cosine corrector. The irradiance of blue light was measured at the sample plane with the LI-190R sensor (LI-COR, USA), combined with the Keithley 6485 picoammeter. Due to the small size of the spot of the red beam at the sample plane in the reflectometric setup, its power was measured with the microscope slide power sensor (S170C, Thorlabs).

### 2.6. Laser scanning confocal microscopy

Microscopic observations were performed with the Axio Observer.Z1 inverted microscope (Carl Zeiss, Germany) and the LSM 880 confocal module. The long-distance LCI Plan-Apochromat 25×, NA 0.8, objective was used with glycerol immersion. Chlorophyll was excited with the 633 nm He–Ne laser, fluorescence emission was recorded in the range of 659 - 721 nm. Chloroplast positioning was observed in the parenchyma of plant leaves or stems irradiated in the reflectometer setup for 1 h. Z-stacks were collected on the adaxial surface of *Draba verna* and *Stellaria media* leaves, the abaxial side of *Tussilago farfara* scale leaves and *Vaccinium myrtillus* stems, with the interval between slices set to ca. 2.4 μm. Projection images were calculated from slices spanning the epidermis and palisade layer (60 - 120 μm of tissue, depending on the species), starting from the upper surface of the epidermis. For cross-sectioning, leaves were fixed with 4% formaldehyde in 50 mM PIPES, 10 mM EGTA, 5 mM MgSO_4_, pH 7.0, and embedded in 6% (w/v) low-melting point agarose (Agarose Super LM, Carl Roth). Leaf cross-sections of 100 – 150 μm in thickness were prepared using the 7000smz^-2^ vibrating microtome (Campden Instruments, UK) with a ceramic blade (7550-1-SS, Campden Instruments). The stems of *V. myrtillus* were cut manually with a razor blade.

### 2.7. X-ray microtomography

For X-ray tomography freshly, collected leaves were cut into small pieces with razor blades and embedded in polyamide (Kapton) tape following a procedure described by (Théroux-Rancourt et al., 2017). X-ray microtomography (µCT) experiments were carried out in parallel beam geometry at the POLYX beamline (Sowa et al., 2023) of the SOLARIS National Synchrotron Radiation Centre (Szlachetko et al., 2023). White X-ray beam from SOLARIS bending magnet was filtered using an Al attenuator with thickness of 0.5 mm and beamline windows (250 µm Be and 150 µm CVD diamond) resulting in a polychromatic X-ray beam with a central energy of approximately 15 keV. Samples on the rotational stage were placed at a distance of 14.5 m from the source. X-ray projections were acquired using a white beam X-ray microscope (Optique Peter, France) with 10 µm or 50 µm thick LuAG:Ce scintillators (Crytur, Czech Republic) and a 10x objective. Sample-to-scintillator distance was set to 40 mm. Images were captured with an sCMOS camera PCO Edge 5.5 with a 6.5 µm pixel size. Tomographic scans were performed over a 180-degree range by a continuous rotation of the sample. For each sample, 2001 equiangular X-ray projections were acquired with a single frame exposure time of 540 ms yielding a total scan time of around 20 minutes. Raw X-ray projections were corrected with flat and dark frames and post-processed with stripe suppression and automatic alignment procedures. Phase retrieval was performed with the Paganin method (Paganin et al., 2002) and δ/β of 100. Tomographic reconstruction was performed using Astra Toolbox (Van Aarle et al., 2016) with an effective voxel size of 0.72 µm. Further segmentation and visualization of tomographic data was performed with a custom-written Mathematica (Wolfram Research, USA) script.

### 2.8. Statistical analysis

The observed distributions of bidirectional reflectance changes substantially deviate from normality when the detector is positioned in such a way as to collect specularly reflected light. The distributions are heavy-tailed, with an excess of extreme values compared to the normal distribution (i.e. high kurtosis). The specular lobe of the leaf reflectance function is relatively sharp, so even relatively small changes in leaf undulations during measurement (*e.g.* induced by the actinic light) may cause part of the specularly reflected beam to miss the detector. In addition, when specularly reflected light is collected, the observed variance depends on the orientation of the polarizer, thus, the homoskedasticity assumption of the standard ANOVA is not satisfied. To reduce the effect of extreme observations, we have transformed the amplitudes and rates of reflectance changes using the inverse hyperbolic sine transformation of the form sinh^-1^(0x)/0, where *x* is the untransformed value of the response variable, while θ is a parameter that needs to be estimated. The linear model of the effect of the plant line, polarizer orientation, and their interaction on the transformed change of reflectance due to actinic light was fitted using the maximum log-likelihood estimator in the *nlme* package (Pinheiro et al., 2023) of the R software (Team R, 2019). The model allowed for heteroskedasticity, with variance free to vary with the orientation of the polarizers. The parameter 0 of the transformation was found through maximization of the concentrated log-likelihood (Burbidge et al., 1988), calculated as the sum of the logarithm of the determinant of the jacobian of the transformation and the maximum log-likelihood for the linear fit. The values of 0 which gave the greatest concentrated log-likelihood were selected. The significance of differences in means of transformed values between groups was assessed with Tukey’s method, using the *emmeans* (Lenth, 2023) and *multcomp* packages (Hothorn et al., 2008). The amplitudes and rates of hemispherical transmittance changes followed the normal distribution, so the statistical analysis was performed on untransformed values. The statistical analysis of transmittance changes was otherwise performed in the same way as for the reflectance measurements.

## 3. Results

### 3.1. The effect of chloroplast movements on leaf reflectance for unpolarized incident light

In this work, we built a goniometric setup to monitor changes in red light leaf reflectance due to blue-light-induced chloroplast movements (Fig. 1). In the first experiment, the setup with one polarizer was used (incident light was unpolarized). To examine the effects of orientation of an *Arabidopsis* leaf on its reflectance and ability to polarize unpolarized incident light, power of *P* and *S-*polarized components of light reflected from the dark-adapted leaves was measured at two angles of incidence and several angles of observation (the angle of observation between 0° and -55° for 40° incidence (Fig. 2 A), and the angle between 15° and - 55° for 55° incidence (Fig. 2 B). In *Arabidopsis* wild-type plants and all mutant lines, *phot1, phot2, phot1phot2, jac1,* and *glabra1,* average bidirectional reflectance ((*P + S*)/2) strongly depended on the observation angle (Fig. 2 and Fig. S2). The average reflectance for wild-type leaves in the direction of specular reflection was ca. 2.2 (40°) or 4.8 (55° incidence) greater than the average reflectance in the direction normal to the leaf surface (Fig. 2). A similar dependence of the bidirectional reflectance on the observation angle was observed for other *Arabidopsis* lines (Fig. S2). Interestingly, reflected radiance was largest for viewing angles exceeding the angle corresponding to the specular reflection from a flat surface. This counterintuitive phenomenon was previously observed for leaves of several species, especially rough ones. It can be explained by a combination of a model of the surface as composed of facets with randomly distributed normal vectors, with Fresnel’s equations of reflectance (Torrance & Sparrow, 1967), (Bousquet et al., 2005). The degree of linear polarization of reflected light varied strongly with the observation angle (Fig. 2 C), especially when the incidence angle was equal to Brewster’s angle.

**Fig. 2.**
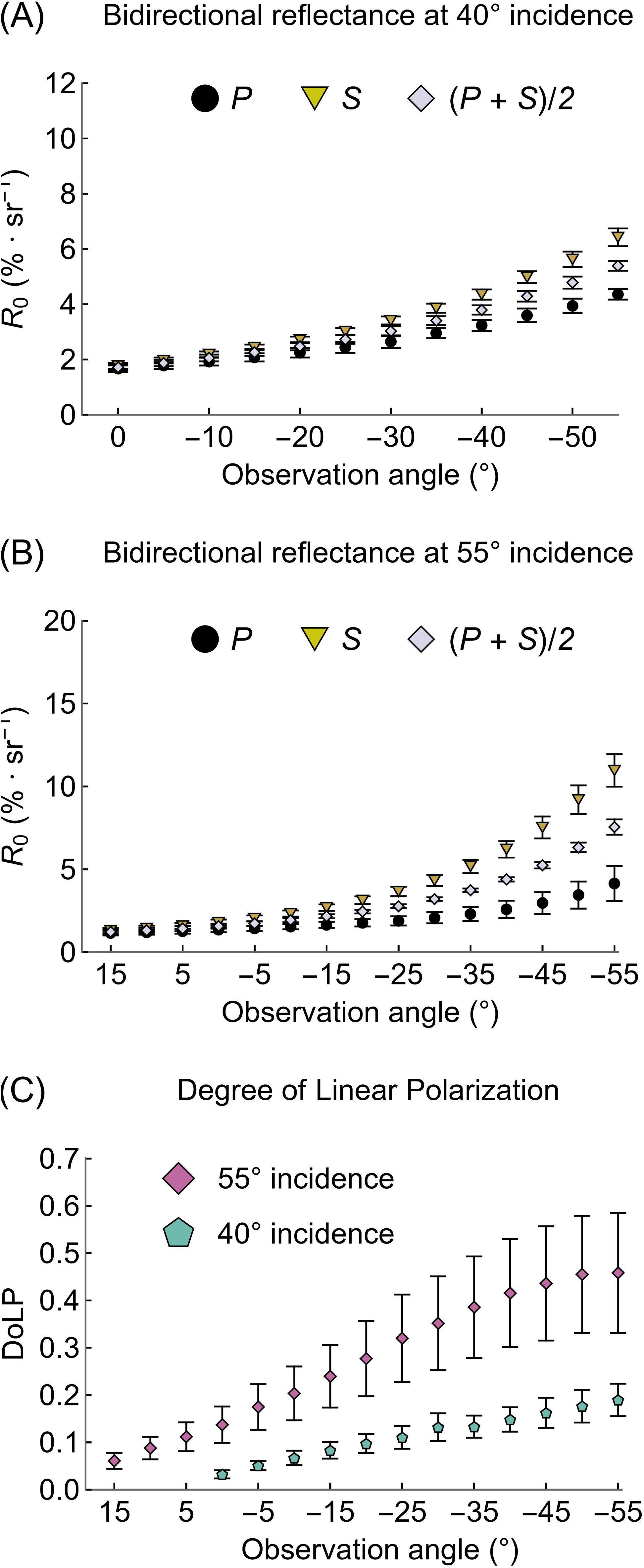
Red-light bidirectional reflectance of the adaxial side of dark-adapted leaves of wild-type *Arabidopsis* plants, measured for the angle incidence of either 40° (A) or 55° (B) and several angles of observation. The incident beam was unpolarized. The transmission axis of the detector polarizer was either parallel (*P,* black dots) or perpendicular (*S*, yellow triangles) to the plane of incidence. Five leaves from different plants were examined for each combination of plant line and angle of incidence. Error bars show the standard error.

To test the accuracy of the reflectance-based monitoring of chloroplast movements, we performed transmittance and reflectance detection in parallel on the same sets of plants. In low blue light of 0.1 µmol·m^-2^·s^-1^, the accumulation response observed in wild-type, *glabra1, phot1,* and *phot2* leaves resulted in a substantial decrease in transmittance (Fig. 3 A, C) and directional reflectance at all combinations of angles examined in this work (40° incidence shown in Fig. 4 A, B, Fig. S3; 55° incidence shown in Fig. 4 E, F, Fig. S4). The changes in reflectance were absent in the *phot1phot2* mutant, which lacks both phototropins. This indicates that the differences observed in wild-type plants were phototropin-dependent. Interestingly, the *jac1* showed a weak, but statistically significant decrease in transmittance at 0.1 µmol·m^-2^·s^-1^ (Fig. 3 A, C). In high blue light of 100 µmol·m^-2^·s^-1^, the avoidance response observed in wild type, *glabra1*, *phot1,* and *jac1* leaves resulted in an increase in transmittance (Fig. 3 B, C) and bidirectional reflectance at both 40° (Fig. 4 C, D, Fig. S5) and 55° (Fig. 4 G, H, Fig. S6) angles of incidence. The effects of the accumulation response observed in *phot2* had the opposite effect. In most cases, no substantial differences in the magnitude of reflectance changes were observed between leaves of wild-type Columbia and the those of the *glabra1* mutant (Fig. 4 A - H), which lack trichomes on their surface. The velocities of chloroplast movements in the analyzed lines monitored through the rates of changes in leaf transmittance d*T/*d*t* and reflectance d*R/*d*t* are shown in Fig. 3 D and Fig. S7.

**Fig. 3.**
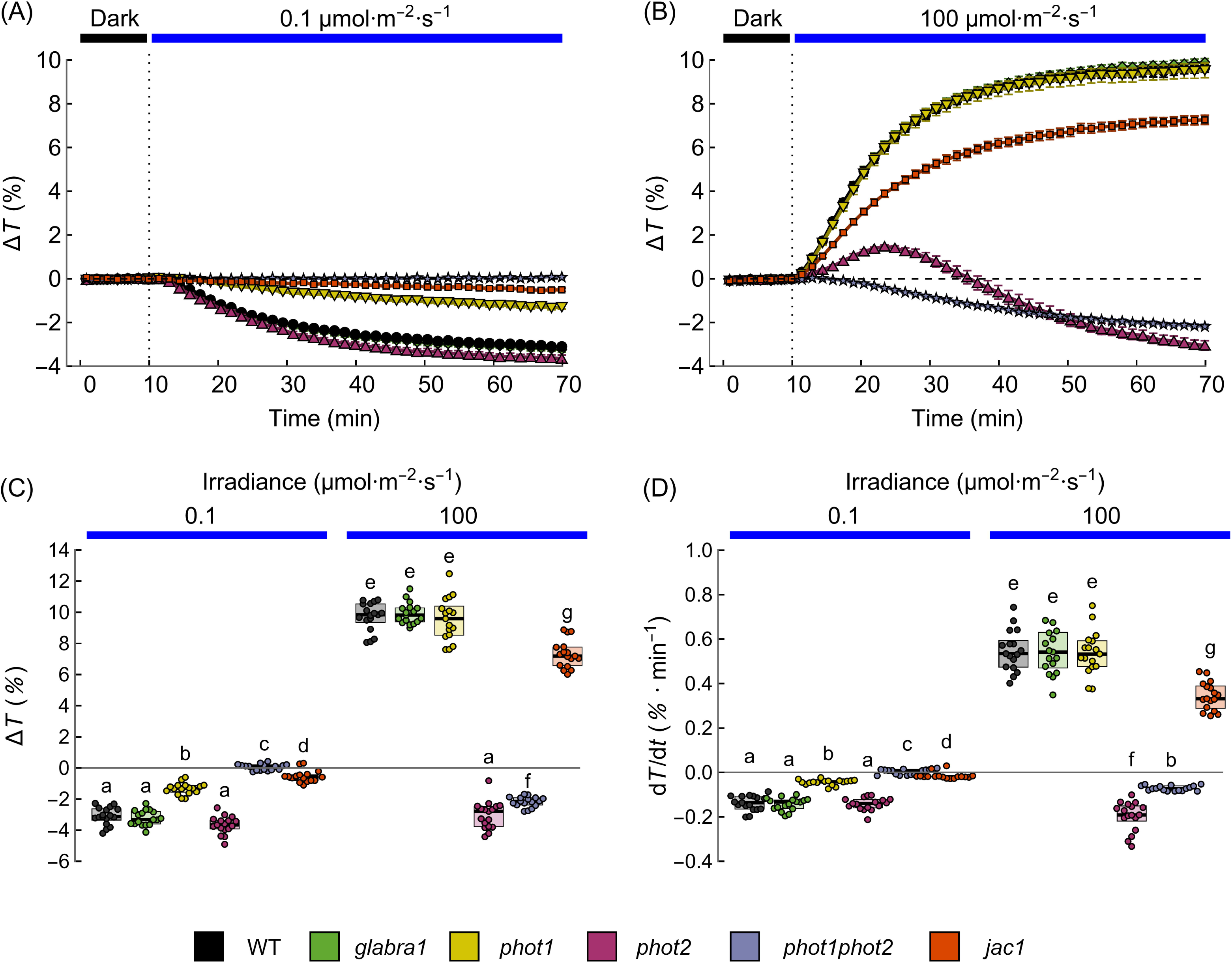
Time courses of changes in hemispherical red light transmittance, induced by blue light of 0.1 µmol·m^-2^·s^-1^ (A) or 100 µmol·m^-2^·s^-1^ (B) in rosette leaves of 5-week-old *Arabidopsis* plants. 17 leaves from different individual plants were measured for each combination of plant line and irradiance level. Error bars show the standard error. Amplitudes (C) and maximal rates (D) of changes in transmittance are calculated from curves in (A) and (B). Boxes mark the interquartile range, horizontal bars across the boxes show the median, and each point marks the value measured for one leaf. The significance of differences between groups is assessed with Tukey’s test on asinh-transformed values. The means of transformed values from groups that do not share a letter are different at the 0.05 level (adjusted for multiple comparisons).

**Fig. 4.**
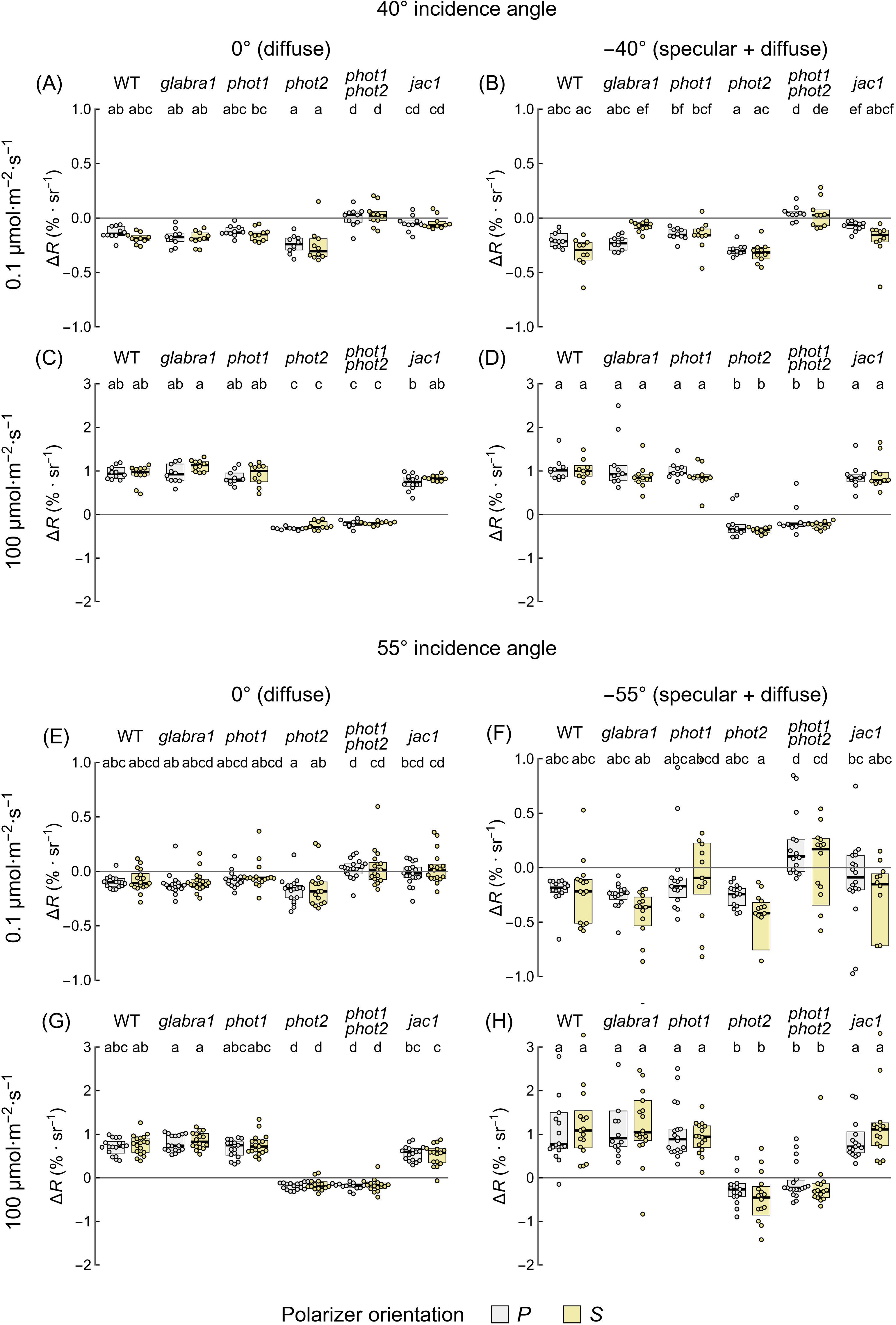
The amplitudes of red-light bidirectional reflectance changes induced by 1 h irradiation with blue light of 0.1 (A, B, E, F) or 100 µmol·m^-2^·s^-1^ (C, D, G, H) in leaves of dark-adapted *Arabidopsis thaliana* plants: wild-type (WT), *glabra1* and chloroplast movement mutants (*phot1*, *phot2*, *phot1phot2*, *jac1*). The blue actinic and red measuring beams were unpolarized and impinged the adaxial leaf surface at the angle of 55° (A-D) or 40° (E-H). For measurements of diffuse directional reflectance (A, C, E, G), the detector was positioned at 0° to the leaf’s normal. For measurements of combined specular and diffuse directional reflectance, the detector was positioned at -55° (B, D) or - 40° (F, H). Two orientations of the detector polarizer were used: its transmission axis was either parallel to the plane of incidence (*P*, attenuates specular reflection, grey boxes) or perpendicular (*S*, yellow boxes). Each dot represents a value obtained for one leaf, with a single leaf harvested from each plant. Boxes show interquartile ranges, while the horizontal bars mark medians. Boxes that do not share any letter represent groups for which the means of transformed values differ at the 0.05 level (Tukey’s method, adjusted for multiple comparisons).

The changes in the power of the *P-*polarized component of reflected light were similar in magnitude to changes in the power of the *S-*polarized component and their means were not statistically different. This was the case for all lines for changes induced both with low (Fig. 4 A, B, E, F) and high light (Fig. 4 C, D, G, H). The variance in the observed reflectance changes was smaller when the polarizer transmitted *P*-polarized light (attenuating specular reflection), as compared to the *S*-orientation. As expected, the initial reflectance *R_0_* of dark-adapted leaves was substantially greater when *S*-polarized light was collected as compared to *P*-polarized light (Fig. S8). Following the approach of (Carter, 1991) and (Baránková et al., 2016), we defined the differential sensitivity of detection as Δ*R/R_0_* (Fig. S9). Such sensitivity was substantially affected by the polarizer orientation when specularly reflected light reached the detector. At Brewster’s angle (55°), the sensitivity of detection for chloroplast avoidance in WT was 2.8 times greater when the polarizer was in the *P* orientation as opposed to the *S* orientation (Fig. S9 H). The absence of any substantial effect of the polarizer orientation on raw values of Δ*R* (Fig. 4) was consistent with the chloroplast positioning affecting only the diffuse reflectance component, originating from the mesophyll.

### 3.2. Effect of chloroplast movements on cross-polarized reflectance and depolarization of polarized light through reflection

To further reduce the specular component and improve the sensitivity of chloroplast movement detection, we inserted an additional polarizer into the path of the measuring beam (Fig. 5). We examined the effects of chloroplast relocations induced by 1 h of 100 µmol·m^-2^·s^-1^ of blue light on reflectance for four combinations of orientations of the polarizers: *P*(incident)/*P*(observed); *P*/*S*; *S*/*P*; *S*/*S*. The experiments were performed on the *Arabidopsis* wild-type and the double *phot1phot2* mutant. The angle of incidence was set to 55°, while the angle of observation was either 0° (diffusely reflected light recorded) or -55° (specularly and diffusely reflected light). The sensitivity Δ*R*/*R*_0_ of detection of the avoidance response in wild-type plants was the highest when polarizers were crossed, i.e. for the *P*/*S* and *S*/*P* combinations. The cross-polarized diffuse (0°observation, Fig. 5 A, E) reflectance increased two times (mean Δ*R*/*R*_0_ ≈ 1) after irradiation. This effect was absent in *phot1phot2* mutant plants that lack chloroplast movements (Fig. 5 C, E), indicating it was indeed due to the avoidance response occurring in the wild type. The small Δ*R*/*R*_0_ values observed when the detector collected specular reflection and both the incident and observed light were *S*-polarized (Fig. 5 B, F) were due to high levels of initial reflectance *R*_0_ (Fig. 5 H), as predicted by Fresnel’s equations. The measured changes of reflectance Δ*R* and the rates d*R/*d*t* are collected in Fig. S 10.

**Fig. 5.**
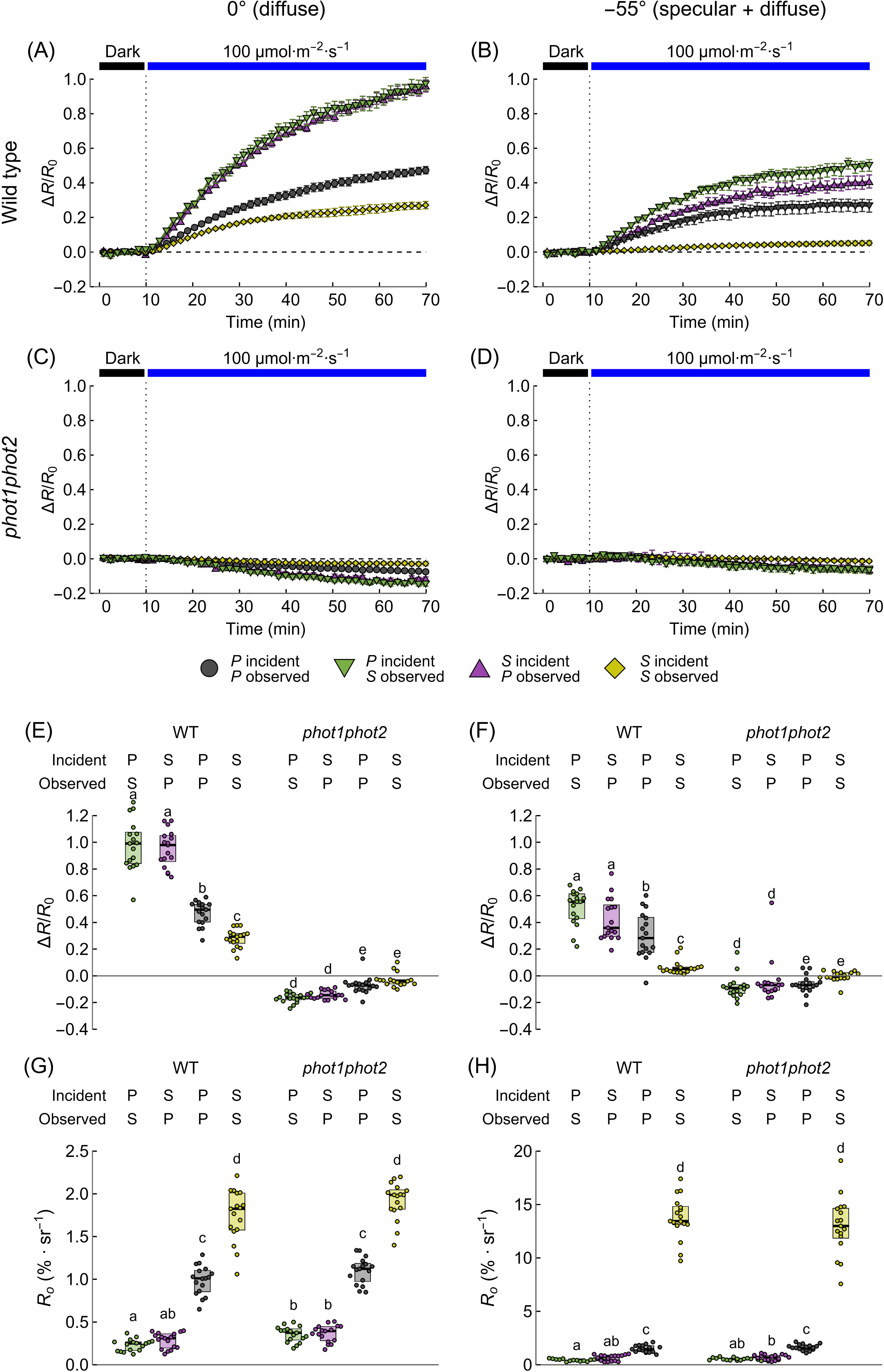
Changes in the polarized red-light bidirectional reflectance induced by 1 h irradiation with high blue light (100 µmol·m^-2^·s^-1^) of the adaxial surface of leaves of wild-type (WT) (A, B) and *phot1phot2* (C, D) *Arabidopsis* plants. Reflectance change curves (A-D) were normalized by the initial reflectance *R_0_* of dark-adapted leaves (G, H). The curves were used to calculate normalized amplitudes Δ*R/R_0_* of reflectance changes (E, F). The blue actinic and red measuring beams impinged the adaxial leaf surface at the angle of 55°. For measurements of diffuse reflectance (A, C, E, G), the detector was positioned at 0° to the leaf normal. For measurements of combined specular and diffuse reflectance (B, D, F, H), the detector was positioned at -55°. Four possible combinations of the orientation (*P/S*) of the polarizers in front of the measuring light source and the detector were examined. Each dot represents a value obtained for one leaf, with a single leaf harvested from each plant. Boxes show interquartile ranges, while the horizontal bars mark medians. Boxes that do not share any letter represent groups for which the means of transformed values differ at the 0.05 level (Tukey’s method, adjusted for multiple comparisons).

We then compared the sensitivity of detection based on cross-polarized (*P* incident, *S* analyzed, 0° observation) reflectance of both accumulation and avoidance responses with that of the transmittance-based detection (Fig. S11). Detached *Arabidopsis* leaves were irradiated for 1 h with 0.1 or 100 µmol·m^-2^·s^-1^ of blue light. To examine unspecific changes in leaf bidirectional reflectance, *e.g.* due to measuring light or leaf dehydration, we also measured the effects of the red light applied without actinic blue light. In *Arabidopsis*, both the avoidance and accumulation responses were easily detected, with mean sensitivities Δ*R*/*R*_0_ of 1.08 (similar to the previous experiment) and -0.23, respectively (Fig. S11 A, B). No substantial change in cross-polarized reflectance was observed when measuring light was applied without the actinic beam. The ratio Δ*R*/*R*_0_ and the analogous sensitivity calculated for hemispherical transmittance, Δ*T*/*T*_0_ (Fig. S11 C, D), were comparable in magnitude and the changes in chloroplast positions were confirmed using microscopic observations (Fig. S11 E).

The observed increase in cross-polarized reflectance may stem from an increase in the leaf’s ability to depolarize linearly polarized light, associated with chloroplast relocations. To test this hypothesis, we examined how chloroplast movements affect the degree of linear polarization (DoLP) of the beam reflected from leaves irradiated with red *P-*polarized light. We performed the measurements of light depolarization for the adaxial and abaxial sides for wild type and *phot1phot2 Arabidopsis* and its changes upon blue light irradiation. The ability of several materials to depolarize light was previously shown to depend on their porosity (Egan et al., 1968). The upper (adaxial) part of the mesophyll in leaves of dicotyledonous plants has the compact form of the palisade parenchyma, characterized by small intercellular spaces. By contrast, the lower (abaxial) part of the mesophyll is occupied by the spongy parenchyma, characterized by high porosity due to large, air-filled spaces (Ma et al., 1990), (Mathers et al., 2018). Differences in the porosity of both types of parenchyma can be seen in the confocal images (Fig. 6 A) and microtomographs (Fig. 6 B) of *Arabidopsis* leaves. Both experimental results and radiative transfer models describing the depolarization of light in a scattering medium (Ma et al., 1990), (Kalshoven et al., 1995) suggest that high chlorophyll content may reduce the leaf’s ability to depolarize light. Wild-type *Arabidopsis* leaves exhibited higher total chlorophyll content than leaves of the phototropin double mutant (Fig. 6 E).

**Fig. 6.**
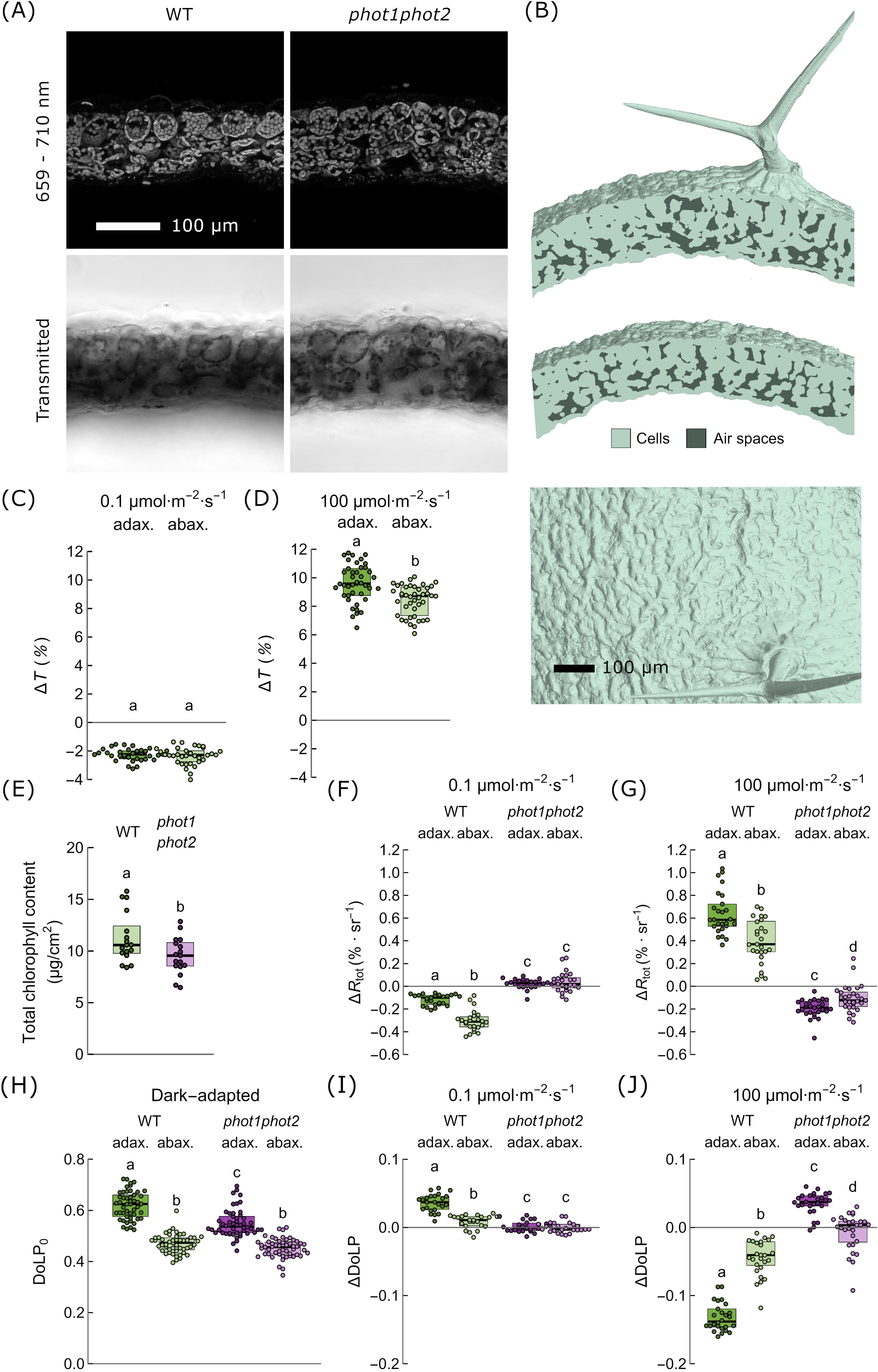
Effect of chloroplast movements on leaf’s ability to depolarize light. (A) Cross-sections of *Arabidopsis thaliana* wild type (WT) and *phot1phot2* mutant leaves imaged with confocal microscopy, using chlorophyll autofluorescence. (B) The adaxial surface and porosity wild-type *Arabidopsis* leaves examined with X-ray microtomography. (C, D) Amplitudes of changes in hemispherical red light transmittance induced by continuous blue light of 0.1 µmol·m^-2^·s^-1^ or 100 µmol·m^-2^·s^-1^ in the abaxial and adaxial side of WT *Arabidopsis* leaves. (E) Total chlorophyll content of *Arabidopsis* WT and *phot1phot2* mutant leaves. (F, G) The amplitude of total polarized red-light bidirectional reflectance Δ*R_tot_* induced by 1 h irradiation with blue light of 0.1 µmol·m^-2^·s^-1^ or 100 µmol·m^-2^·s^-1^ of the adaxial and abaxial surfaces of wild-type (WT) and *phot1phot2 Arabidopsis* leaves. (H-J) The degree of linear polarization (DoLP) of the adaxial and abaxial surfaces of wild-type (WT) and *phot1phot2 Arabidopsis* leaves in darkness (H) or after 1 h irradiation with blue light of 0.1 µmol·m^-2^·s^-1^ (I) or 100 µmol·m^-2^·s^-1^ (J). In (C-J) each dot represents a value obtained for one leaf, with a single leaf harvested from each plant. Boxes show interquartile ranges, while the horizontal bars mark medians. Boxes that do not share any letter represent groups for which the means of transformed values differ at the 0.05 level (Tukey’s method, adjusted for multiple comparisons).

Chloroplast movements can be induced through irradiation of both the adaxial and abaxial sides of the leaf. In wild-type *Arabidopsis*, we observed similar decreases in hemispherical transmittance Δ*T* of the leaf when abaxial and adaxial sides were irradiated with low blue light of 0.1 µmol·m^-2^·s^-1^ (Fig. 6 C). High blue light of 100 µmol·m^-2^·s^-1^ resulted in a greater increase in transmittance when the adaxial side was irradiated (Fig. 6 D). The measurements of the effects of chloroplast movements on the leaf’s ability to depolarize light were performed for red *P*-polarized light at 55° incidence. We measured the power of *P* and *S* components of reflected light at 0° observation angle, before and after irradiation with low (0.1 µmol·m^-2^·s^-1^) (Fig. S12 A-D) or high (100 µmol·m^-2^·s^-1^) blue light (Fig. S12 E-H). Then, we calculated how the total leaf reflectance *R_tot_* (Fig. 6 F, G) and the degree of linear polarization DoLP (Fig. 6 I, J) of reflected red light changed after irradiation with blue light. In wild-type plants, Δ*R_tot_*decreased after irradiation with low blue light and increased after irradiation with high blue light regardless of the leaf side. This effect was not observed for the *phot1phot2* mutant (Fig. 6 F, G). For dark-adapted leaves, light reflected from the adaxial side retained a greater degree of polarization than light reflected from the abaxial side. This effect was observed for both the wild type and the double phototropin mutant (Fig. 6 H). For wild-type plants, red light retained a higher degree of linear polarization upon reflection from low blue-light irradiated leaves as compared to dark-adapted ones (ΔDoLP > 0) (Fig. 6 I). This effect was absent in *phot1phot2* mutant, which suggests that the reduced ability of leaves to depolarize incident *P-*polarized red light after low-blue-light irradiation was due to chloroplast accumulation. By contrast, the degree of polarization of red light reflected from wild-type leaves decreased after irradiation with high blue light (ΔDoLP < 0) (Fig. 6 J). No drop in DoLP of reflected light was observed for the *phot1phot2* mutant, suggesting that the effect observed in wild-type leaves was due to the chloroplast avoidance response. Irradiation of the adaxial side of *phot1phot2* leaves with high blue light had the opposite effect to that observed for the wild type. An increase in DoLP of the reflected red beam was observed, consistent with a small decrease of leaf transmittance (Fig. 3 B, C).

### 3.3. Chloroplast movements in species collected in the natural environment

To assess the potential applicability of cross-polarized reflectance for the detection of chloroplast relocations in field studies, we examined detached photosynthetic organs of four wild species, irradiated with 0 (measuring beam only), 0.1 or 100 µmol·m^-2^·s^-1^ of blue light for 1 h. Measurements were recorded on rosette leaves of *Draba verna* (Fig. 7 A), a close relative of *Arabidopsis*, stem leaves of *Stellaria media* (Fig. 7 B), scales covering flowering shoots of *Tussilago farfara* (Fig. 8 A), and green stems of *Vaccinium myrtillus* (Fig. 8 B). The red measuring beam did not induce substantial changes in reflectance or transmittance in any of the investigated species. Pronounced chloroplast accumulation and avoidance responses were observed through microscopic observations in *D. verna* (Fig. 7 A) and *S. media* (Fig. 7 B) leaves. As in *Arabidopsis*, the accumulation response led to a substantial decrease in cross-polarized reflectance and transmittance in both species, while the avoidance response had the opposite effect. In the scales of *T. farfara*, only the avoidance response to high blue light could be observed microscopically. It was accompanied by an easily detectable increase in reflectance (mean Δ*R*/*R*_0_ of 0.24) and transmittance (Fig. 8 A). A small, but statistically significant decrease in transmittance and reflectance, corresponding to chloroplast accumulation, was observed after low light irradiation. We also investigated chloroplast relocations in green, photosynthetic stems of *Vaccinium myrtillus,* collected in early spring, at the beginning of bud break (Fig. 8 B). While the transmittance-based method could not be used for opaque stems, the sizeable (mean Δ*R*/*R*_0_ of 0.31) changes in cross-polarized reflectance induced by irradiation with high blue light pointed to the chloroplast avoidance response. Low blue light led to a decrease in reflectance (mean Δ*R*/*R*_0_ of -0.05).

**Fig. 7.**
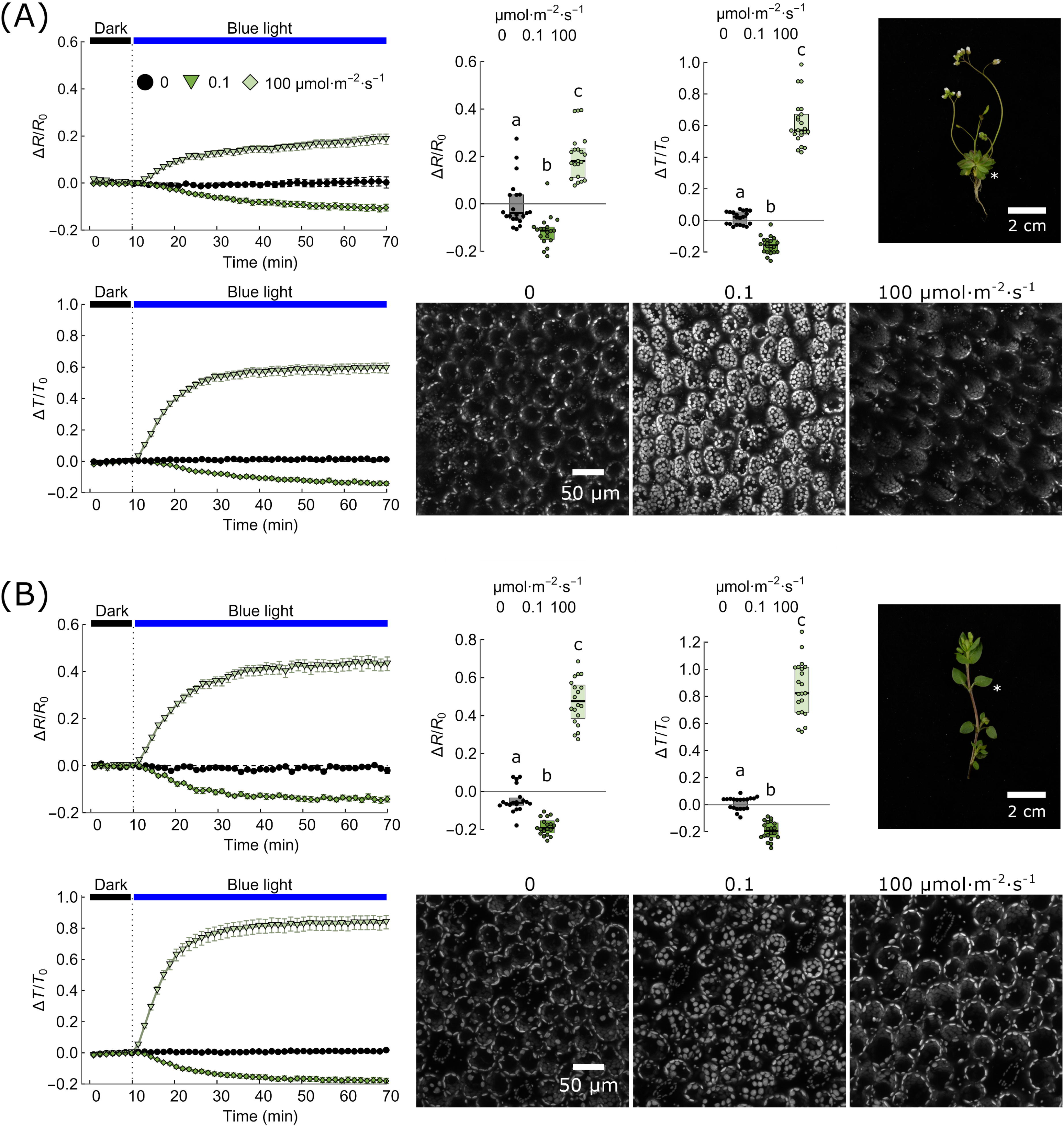
Chloroplast movements examined with photometric, reflectometric, and microscopic observations *Draba verna* (A) and *Stellaria media* (B) leaves. Time courses of changes in red-light bidirectional reflectance and hemispherical transmittance induced by 0.1 or 100 µmol·m^-2^·s^-1^ of blue light or mock irradiation of the adaxial side of leaves. The angle of incidence of beams was 55° while the angle of observation was 0° (diffuse reflectance recorded). The incident, measuring light was polarized in the direction parallel to the plane of incidence (*P*). The polarizer in front of the detector (analyzer) transmitted light perpendicular to the incidence plane (*S*). Each dot represents a value obtained for one leaf harvested from a different plant. Boxes show interquartile ranges, while the horizontal bars mark medians. The means of transformed values from groups that do not share any letter differ at the 0.05 level (Tukey’s method, adjusted for multiple comparisons). Confocal microscopy images of leaf tissue were recorded after 1 h irradiation with blue light of 0.1 µmol·m^-2^·s^-1^ or 100 µmol·m^-2^·s^-1^ or mock irradiated. Chlorophyll autofluorescence was detected. Photographs show the developmental stage of plants at the time of the measurement, with the examined leaves marked with asterisks.

**Fig. 8.**
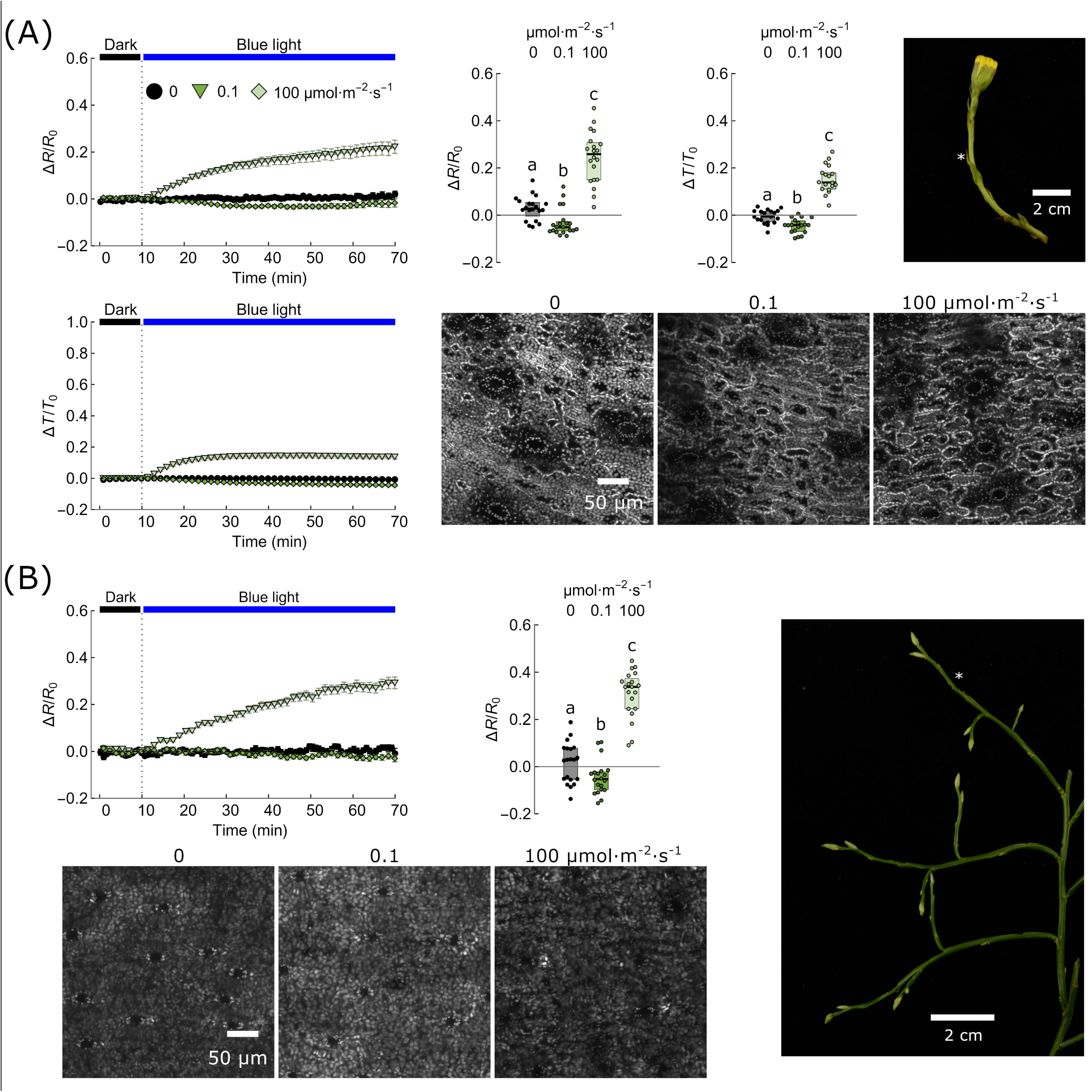
Chloroplast movements examined with photometric, reflectometric, and microscopic observations *Tussilago farfara* (A) and *Vaccinium myrtillus* (B). Time courses of changes in red-light bidirectional reflectance and hemispherical transmittance induced by 0.1 or 100 µmol·m^-2^·s^-1^ of blue light or mock irradiation of the sun-exposed (abaxial) *Tussilago* scales or *Vaccinium* stems. The angle of incidence of beams was 55° while the angle of observation was 0° (diffuse reflectance recorded). The incident, measuring light was polarized in the direction parallel to the plane of incidence (*P*). The polarizer in front of the detector (analyzer) transmitted light perpendicular to the incidence plane (*S*). Each dot represents a value obtained for one leaf harvested from a different plant. Boxes show interquartile ranges, while the horizontal bars mark medians. The means of transformed values from groups that do not share any letter differ at the 0.05 level (Tukey’s method, adjusted for multiple comparisons). Confocal microscopy images of *Tussilago* scales and *Vaccinium* stems were recorded after 1 h irradiation with blue light of 0.1 µmol·m^-2^·s^-1^ or 100 µmol·m^-2^·s^-1^ or mock irradiated. Chlorophyll autofluorescence was detected. Photographs show the developmental stage of plants at the time of the measurement, with the examined organs marked with asterisks.

To link the changes in optical properties with the structure of the leaf, we investigated cross-sections of leaves with confocal microscopy (Fig. S13) and larger leaf fragments with X-ray microtomography (Fig. 9). The leaves of *D. verna* were ca. 200 – 250 µm thick, consisting of 2-3 layers of palisade mesophyll and 3-4 layers of spongy mesophyll (Fig. S13 A, Fig. 9. A). The upper epidermis was covered with trichomes similar in shape to those of *Arabidopsis* (Fig. 9 A). The leaves of *S. media* consisted of only 1-2 layers of palisade mesophyll, characteristic of species growing in shaded environments (Fig. S13 B, Fig. 9 B). The extracellular spaces occupied a high fraction of the leaf (Fig. 9 B). The epidermis comprised pavement cells with lobes and indentations (Fig. 9 B). Epidermal cells of *D. verna* and *S. media* do not show any preferred direction in their shape or orientation. By contrast, the thin, monofacial scales of *T. farfara* were covered with epidermal cells elongated along a single axis (Fig. S13 C, Fig. 9 C). Thus, any rotation of the scale around the normal to the blade may affect specular reflection off its surface. *V. myrtillus* stem was rectangular on its cross-section, with clear corners. It consisted of two types of cells containing chloroplasts (Fig. S13 D). 2-3 layers of small cells rich in chloroplasts localized just under the epidermis. Larger cells that form loose parenchyma with intracellular spaces fill the inner part of the stem (Fig. S13 D).

**Fig. 9.**
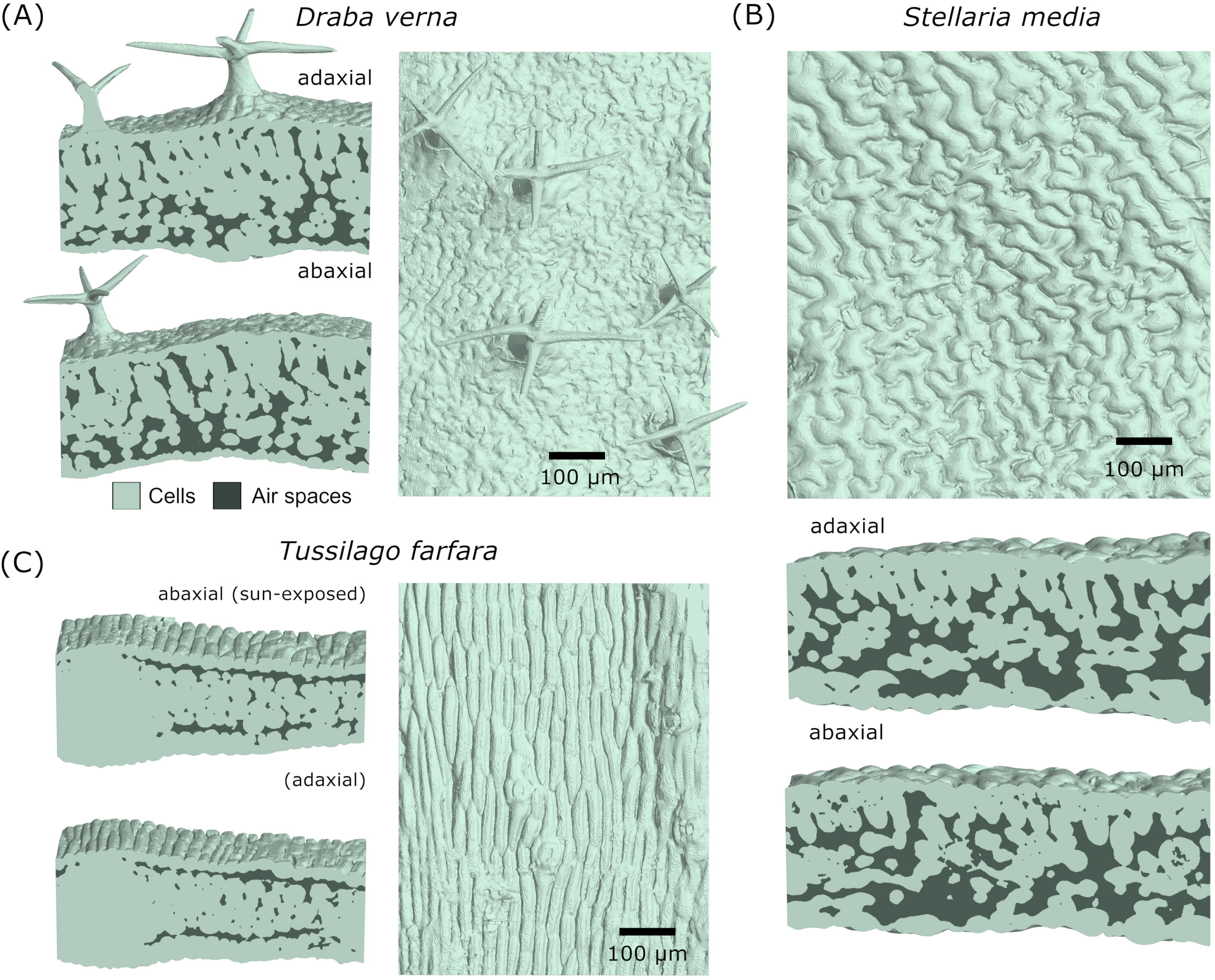
Leaf microtomography images of *Draba verna* (A)*, Stellaria media* (B)*, Tussilago farfara* (C). For each species, a leaf cross-section and the sun-exposed surface are shown. Scale bar 100 µm.

## 4. Discussion

We show that changes in leaf directional reflectance mimic the changes in hemispherical transmittance, monitored in the standard, photometric approach for chloroplast movement detection, in agreement with previous reports (Dutta et al., 2015), (Seybold, 1955) (Zurzycki, 1961). Using a set of chloroplast movement mutants of the model plant *Arabidopsis thaliana*, we demonstrate that the observed changes in reflectance stem from different positions of chloroplasts, not from other light-dependent processes that occur in the leaf (Fig. 4). In the examined wild plant species, the relative sensitivity of reflectance-based detection was close to the sensitivity of the transmittance-based method, suggesting that remote monitoring of chloroplast movements in field conditions may be feasible (Fig. 7 - 8).

Our results indicate that chloroplast movements affect the diffuse component of reflectance (Fig. 4), while its specular component does not carry any information about chloroplast positioning. This is in line with the fact that specular reflection is not affected by the pigment composition of the leaf, as shown in studies on variegated leaves (Grant et al., 1987). Specular reflection arises mainly from the interaction between incoming radiation and the waxes or the cuticle. The prominence of specular lobes of BRDF depends on the angle of incidence and the roughness of the leaf surface (Breece & Holmes, 1971), (Bousquet et al., 2005). In our experiments, the presence of trichomes does not exert a considerable effect on light reflected from the leaves of *Arabidopsis thaliana* (hairy WT Col-0 versus bare *gl1*) (Fig. 4). This is consistent with previous results obtained for the *glabra1* mutant showing no significant difference in specular reflectance between this hairless line and lines bearing trichomes (Neuwirthová et al., 2021).

Strong dependence of detected leaf reflectance on incidence and observation angles may hinder the detection procedure (Fig. 4). Leaf movements (*e.g.* nyctinastic or due to wind) may increase the variance of reflectance signal, but the deviation can be expected to average out when a considerable number of measurements is taken. Systematic errors, however, can arise from light-induced reorientation of leaves and shoots, either photonastic, phototropic, or due to wilting. Light-induced changes in leaf orientation have been observed in several plant families. They can be fast and substantial in magnitude. In *Oxalis oregana*, high blue light induces leaflet folding, with the petiole angle decreasing by 90° within a few minutes (Björkman & Powles, 1981). In *Arabidopsis*, the blade of a leaf changes its orientation within a few hours in response to low blue light (Inoue et al., 2008). Such a rotation of the blade would change the angle of incidence of measuring light, affecting the amount of reflected light received by the detector. The dependence of reflectance of an *Arabidopsis* leaf on the observation angle is pronounced (Fig. 2), even though leaves in this species are not particularly glossy. This suggests that a considerable magnitude of the spurious signal can be expected if chloroplast movements are examined through reflectance measurements of leaves that move, *e.g.* in plants exposed to wind.

We show that this issue can be mitigated through polarization-sensitive detection (Fig. 5). This approach improves the relative sensitivity of detection and reduces the angular dependence of the magnitude of the changes in leaf reflectance. Our results imply that cross-polarized reflectance may be used to investigate chloroplast movements in several species growing outdoors which largely differ in leaf morphology and architecture (Fig. 7, 9). It is also a convenient method for detecting chloroplast avoidance in photosynthetic stem tissues (Fig. 8 B). Green stems of *V. myrtillus* can carry out photosynthesis even in winter and early spring when the shoots are leafless and covered with snow (Havas, 1971). They are equipped with stomata and photosynthetic tissue located directly beneath the epidermis, a feature shared with several genera of shrubs inhabiting extreme environments (Nilsen, 1995), (Ávila et al., 2014). Studies in Norwegian tundra indicate that stem photosynthesis corresponds to ca. 10 – 15% of leaf photosynthesis in summer (Skre, 1975). In deciduous shrubs, such as *V. myrtillus*, stem photosynthesis may play an important role in the fall and early spring, when temperatures are low. In such conditions, the sensitivity of chloroplast movements to light in leaves increases (Łabuz et al., 2015).

The advantages of remote sensing of chloroplast movements would be twofold. The first advantage is practical, as chloroplast movements influence not only the total reflectance of the leaf but also the shape of the spectrum of reflected light. As a result, they may affect values of vegetation indices used in remote sensing, in particular indices related to the pigment content (Hermanowicz & Łabuz, 2024). Thus, a reliable method of detection of chloroplast positioning in conditions in which the leaf orientation or canopy geometry cannot be controlled may allow for the reduction of a confounding factor in the reflectance-based detection of other plant traits. In addition, reflectance-based detection appears to be the only method that can achieve the throughput necessary to investigate the ecological role of chloroplast movements in ecosystems. The variability of chloroplast movements between different species is high (Königer & Bollinger, 2012). In addition, they exhibit substantial phenotypic plasticity, with their magnitude dependent on the light conditions during growth (Trojan & Gabrys, 1996). Thus, direct methods of chloroplast movement detection, through microscopic observations or transmittance measurements, appear too labor-intensive to provide a full picture of the significance of this process in the field conditions. The substantial effect of the chloroplast distribution on the cross-polarized reflectance and depolarization of light upon reflection from the leaf seems to indicate that it might be possible to detect it using techniques of polarimetric remote sensing, for example, LiDAR-based Mueller matrix polarimetry (Keyser et al., 2019).

## 5. Conclusions

The effect of chloroplast movements on directional leaf reflectance appears to be substantial, however, the estimation of the magnitude of response requires care. Changes in the signal due to light-induced leaf reorientation may be then mistaken for chloroplast relocations. In this work, we show that measurements of cross-polarized reflectance alleviate this problem, enabling sensitive non-contact detection of chloroplast movements. The developed methodology may be combined with already available hardware for remote sensing to improve the assessment of physiological traits in the field.

## Acknowledgments

We would like to thank Anna Kozłowska-Mroczek for her excellent assistance in preparing the plant material for the measurements. We thank Mariusz Mroczek and Aneta Prochwicz for their technical support. We thank Tomasz Kołodziej and Paweł Wróbel for help at the POLYX beamline of the SOLARIS National Synchrotron Radiation Centre.

## Author contributions

P.H. - Conceptualization; Data curation; Formal analysis; Funding acquisition; Investigation; Methodology; Resources; Software; Validation; Visualization; Writing – original draft; review & editing, A.G. - Investigation, K.S. - Resources; Software, P.K. Data analysis, J.L. - Conceptualization; Investigation; Resources; Supervision; Writing – original draft, review & editing.

## Data availability

The data that support the findings of this study are openly available in FigShare at https://doi.org/10.6084/m9.figshare.21082654 (reflectance and transmittance recordings for *Arabidopsis* WT and mutants) and https://doi.org/10.6084/m9.figshare.24424843 (wild plants). Java source code for the software is publicly available via GitHub at https://github.com/pawelHerm/beamJ/tree/master/BeamJ. Wolfram Mathematica package for ray tracing is available at https://github.com/plantPhotobiologyLab/openRayTracer.

