## Supporting material for "Sensitive detection of chloroplast movements through changes in leaf cross-polarized reflectance"

**Supplementary figures**

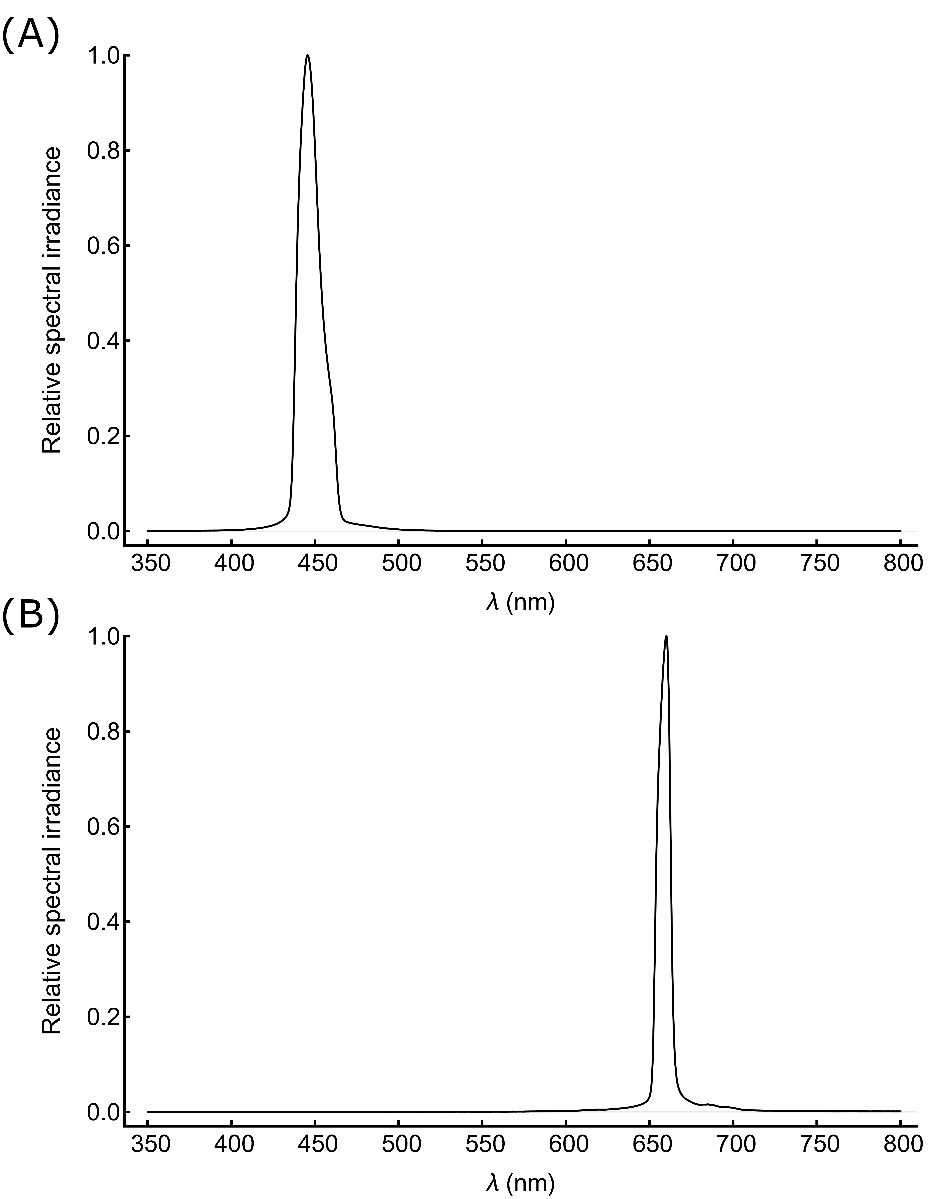

Fig. S1. Relative spectral irradiance as the function of wavelength of the (A) actinic (measured maximum at 446 nm, FWHM 15 nm) and (B) measuring (660 nm, FWHM 9 nm) beams used in the reflectometric setup. The spectra were recorded with the Black Comet SR spectrometer (Stellar-Net). The cosine corrector of the probe was positioned in the leaf sample mount and oriented perpendicularly to the beams.

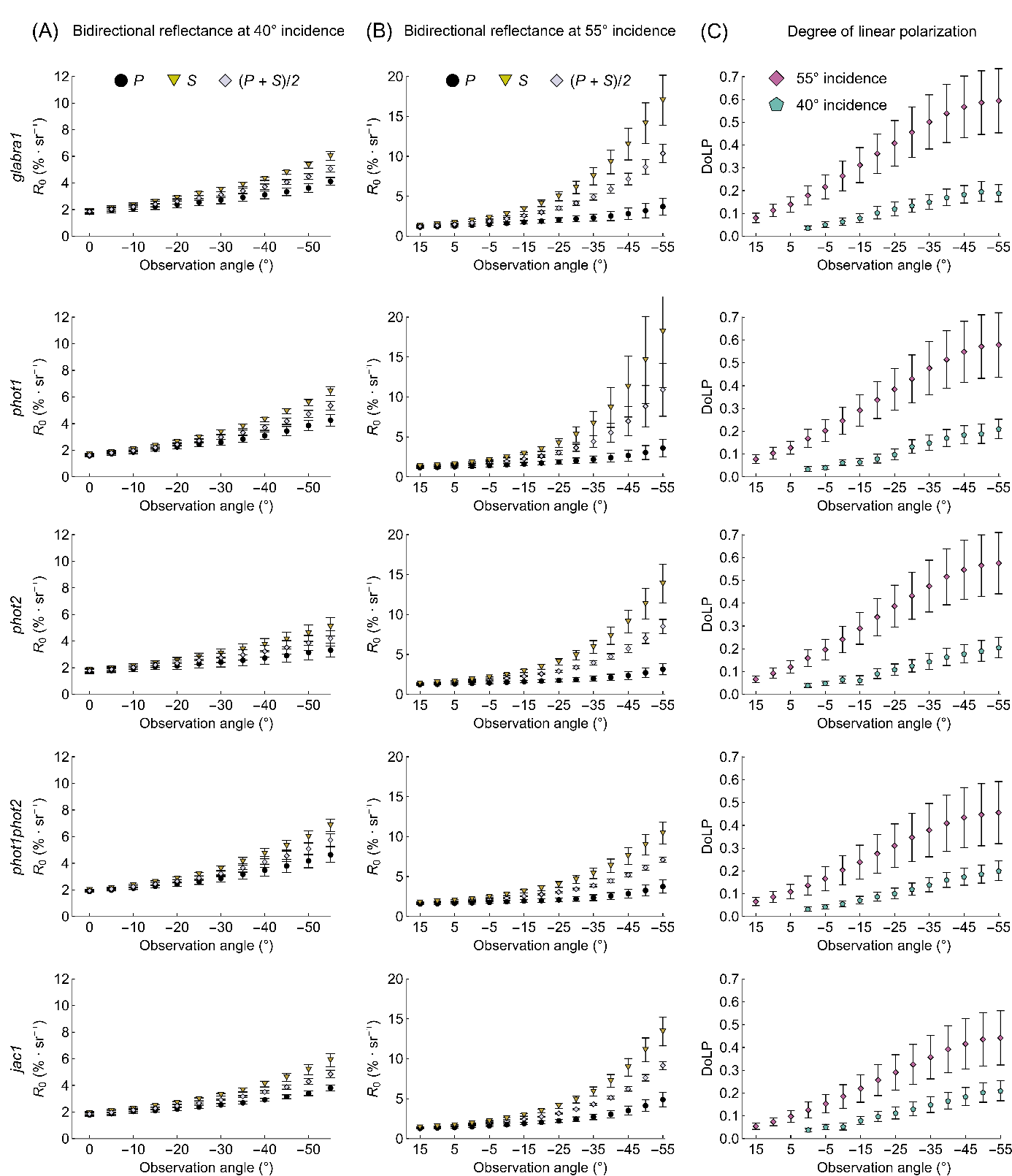

Fig. S2. Red-light bidirectional reflectance of the adaxial side of dark-adapted leaves of *glabra1*, *phot1*, *phot2*, *phot1phot2,* and *jac1* *Arabidopsis* plants, measured for the angle incidence of either 40° (A) or 55° (B) and several angles of observation. The incident beam was unpolarized. The transmission axis of the detector polarizer was either parallel (*P,* black dots) or perpendicular (*S*, yellow triangles) to the plane of incidence. The degree of linear polarization (DoLP) of reflected light (C) was estimated from the data in (A, B) as (*R_s_* - *R_P_*)/(*R_s_* + *R*_P_), where *R_s_* and *R_P_* are reflectance values obtained with the *S* and *P* orientations of the polarizer, respectively. Five leaves from different plants were examined for each combination of plant line and angle of incidence. Error bars show the standard error.

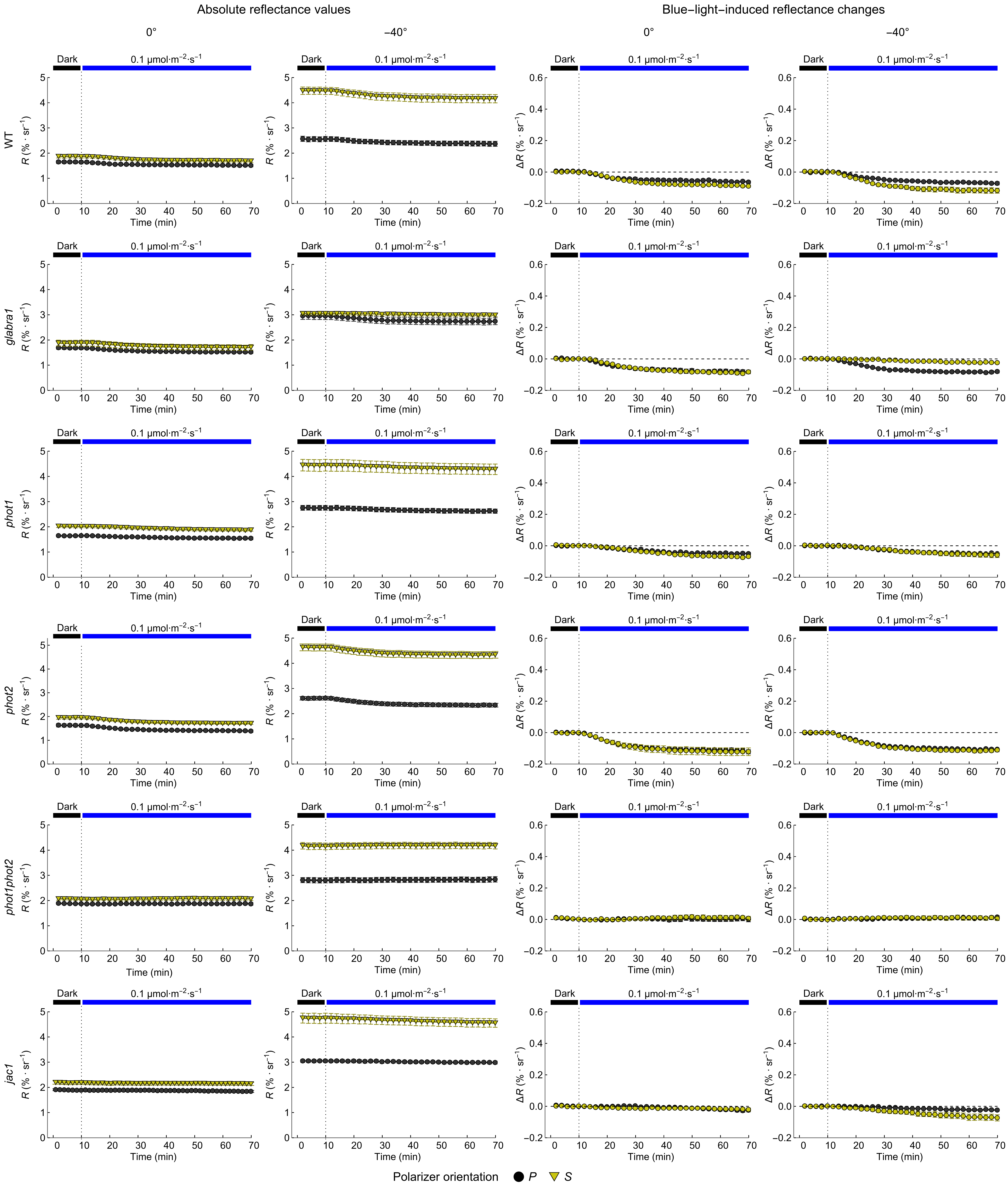
 Fig. S3. Time courses of changes in red-light bidirectional reflectance induced by 1 h irradiation with low blue light (0.1 µmol·m^-2^·s^-1^) of the adaxial surface of leaves of *Arabidopsis* plants. As marked in the figure, each row of the matrix of charts corresponds to a different plant line, from the top: wild type (WT), *glabra1, phot1*, *phot2*, *phot1phot2,* and *jac1* mutants. The angle of incidence of the measuring and actinic beams was 40°. The incident beams were not polarized. The detector was positioned at the angle of 0° (the detector normal parallel to the leaf normal) or -40°. The transmission axis of the detector polarizer was either parallel (*P,* black dots) or perpendicular (*S*, yellow triangles) to the plane of incidence. The charts in the first two columns show raw reflectance recordings *R*, while the curves in the third and fourth columns show the difference between current reflectance and the reflectance at the onset of blue light irradiation *ΔR*. Error bars show the standard error, the plot markers (black dots for the *P* polarizer orientation, yellow triangles for the *S* orientation) show mean values.

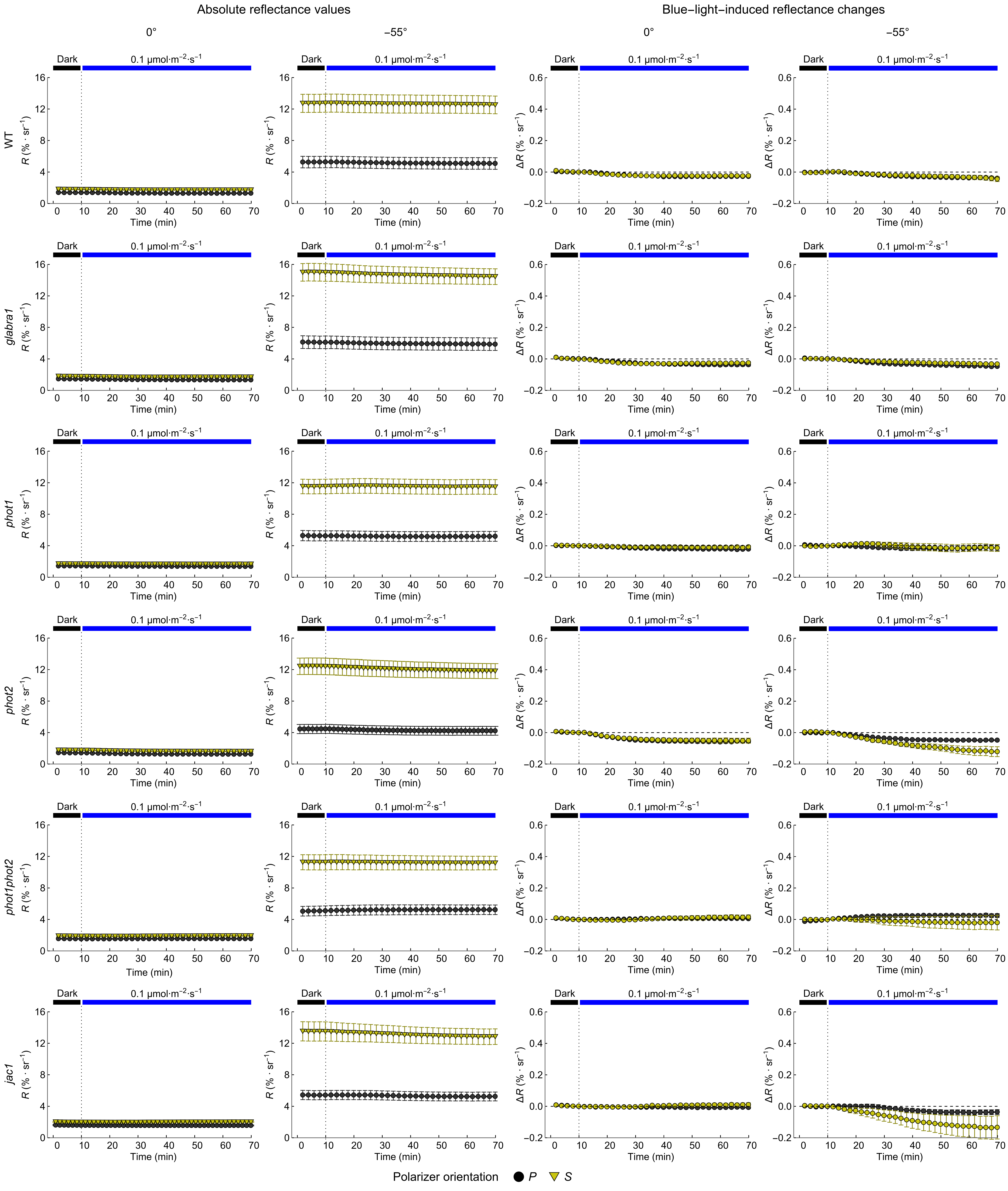

Fig. S4. Time courses of changes in red-light bidirectional reflectance induced by 1 h irradiation with low blue light (0.1 µmol·m^-2^·s^-1^) of the adaxial surface of leaves of *Arabidopsis* plants. As marked in the figure, each row of the matrix of charts corresponds to a different plant line, from the top: wild type (WT), *glabra1, phot1*, *phot2*, *phot1phot2,* and *jac1* mutants. The angle of incidence of the measuring and actinic beams was 55°. The incident beams were not polarized. The detector was positioned at the angle of 0° (the detector normal parallel to the leaf normal) or -55°. The transmission axis of the detector polarizer was either parallel (*P,* black dots) or perpendicular (*S*, yellow triangles) to the plane of incidence. The charts in the first two columns show raw bidirectional reflectance recordings *R*, while the curves in the third and fourth columns show the difference *ΔR* between current reflectance and the reflectance at the onset of blue light irradiation. Error bars show the standard error. The plot markers (black dots for the *P* polarizer orientation, and yellow triangles for the *S* orientation) show mean values.

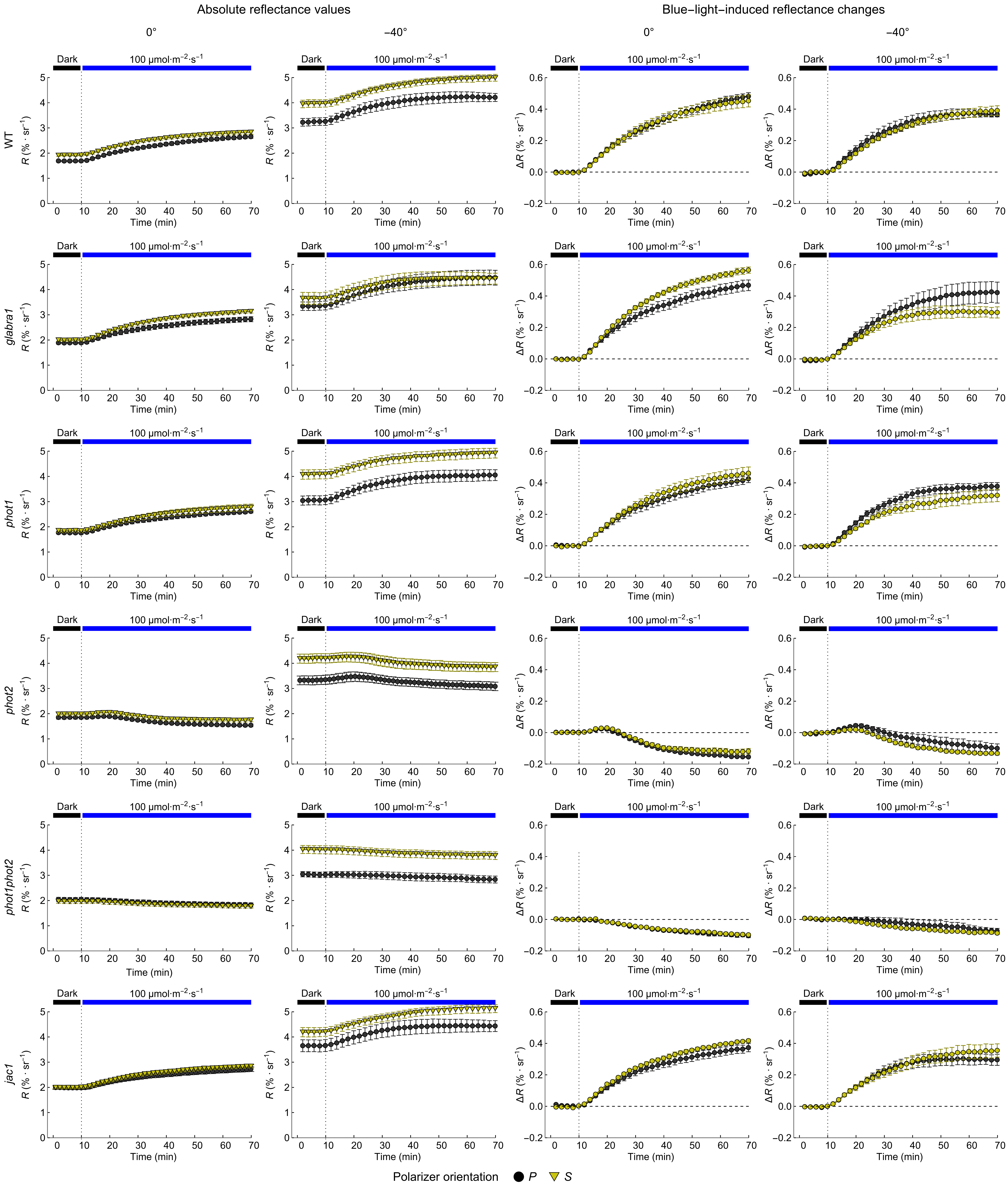
 Fig. S5. Time courses of changes in red-light bidirectional reflectance induced by 1 h irradiation with high blue light (100 µmol·m^-2^·s^-1^) of the adaxial surface of leaves of *Arabidopsis* plants. As marked in the figure, each row of the matrix of charts corresponds to a different plant line, from the top: wild type (WT), *glabra1, phot1*, *phot2*, *phot1phot2,* and *jac1* mutants. The angle of incidence of the measuring and actinic beams was 40°. The incident beams were not polarized. The detector was positioned at the polar angle of 0° (the detector normal parallel to the leaf normal) or -40°. The transmission axis of the detector polarizer was either parallel (*P,* black dots) or perpendicular (*S*, yellow triangles) to the plane of incidence. The charts in the first two columns show raw reflectance recordings *R*, while the curves in the third and fourth columns show the difference between current reflectance and the reflectance at the onset of blue light irradiation *ΔR*. Error bars show the standard error, the plot markers (black dots for the *P* polarizer orientation, yellow triangles for the *S* orientation) show mean values.

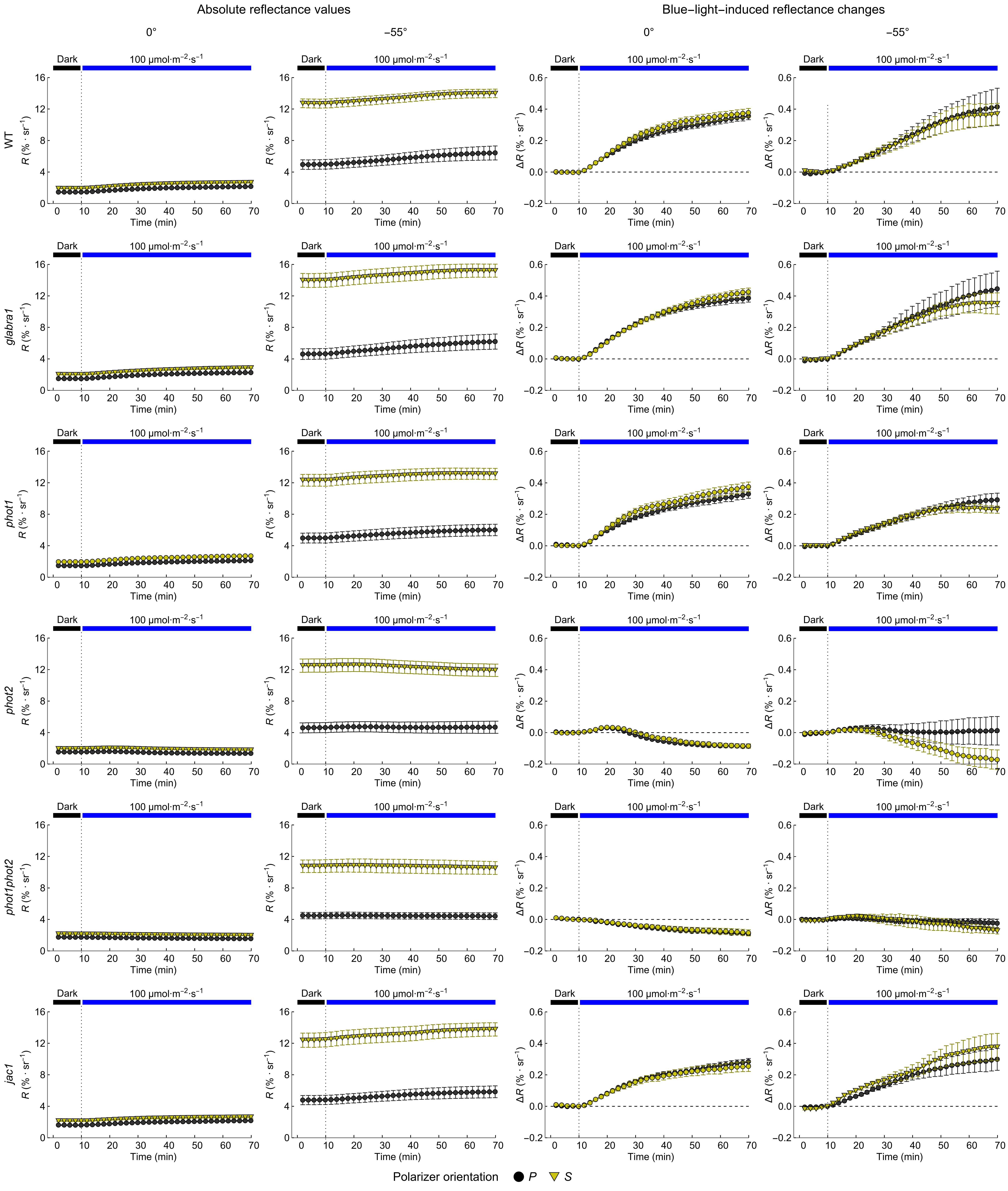
Fig. S6. Time courses of changes in red-light bidirectional reflectance induced by 1 h irradiation with high blue light (100 µmol·m^-2^·s^-1^) of the adaxial surface of leaves of *Arabidopsis* plants. As marked in the figure, each row of the matrix of charts corresponds to a different plant line, from the top: wild type (WT), *glabra1, phot1*, *phot2*, *phot1phot2,* and *jac1* mutants. The angle of incidence of the measuring and actinic beams was 55°. The incident beams were not polarized. The detector was positioned at the angle of 0° (the detector normal parallel to the leaf normal) or -55°. The transmission axis of the detector polarizer was either parallel (*P,* black dots) or perpendicular (*S*, yellow triangles) to the plane of incidence. The charts in the first two columns show raw reflectance recordings *R*, while the curves in the third and fourth columns show the difference between current reflectance and the reflectance at the onset of blue light irradiation *ΔR*. Error bars show the standard error. The plot markers (black dots for the *P* polarizer orientation, and yellow triangles for the *S* orientation) show mean values.

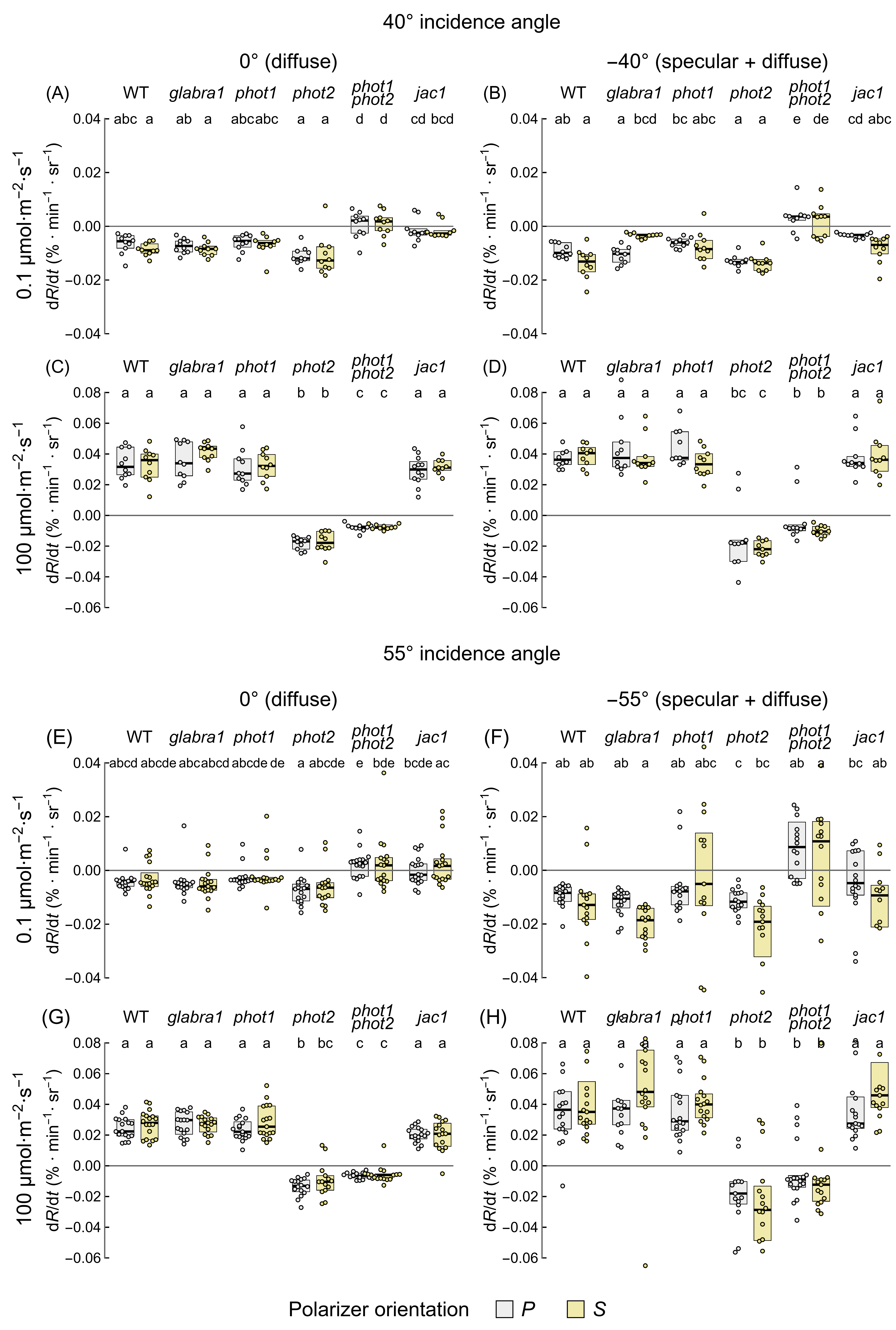

Fig. S7. The maximal rates of red-light bidirectional reflectance changes induced by 1 h irradiation with blue light of 0.1 (A, B, E, F) or 100 µmol·m^-2^·s^-1^ (C, D, G, H) in leaves of dark-adapted *Arabidopsis thaliana* plants: wild type (WT), *glabra1* and chloroplast movement mutants (*phot1*, *phot2*, *phot1phot2*, *jac1*). The blue actinic and red measuring beams were unpolarized and impinged the adaxial leaf surface at the angle of 40° (A-D) or 55° (E-H). For measurements of diffuse directional reflectance (A, C, E, G), the detector was positioned at 0° to the leaf normal. For measurements of combined specular and diffuse directional reflectance, the detector was positioned at -40° (B, D) or -55° (F, H). For each observation angle, two orientations of the detector polarizer were used: its transmission axis was either parallel to the plane of incidence (*P*, attenuates specular reflection, grey boxes) or perpendicular (*S*, yellow boxes). Each dot represents a value obtained for one leaf, with a single leaf harvested from each plant. Boxes show interquartile ranges, while the horizontal bars mark medians. Boxes that do not share any letter represent groups for which the means of transformed values differ at the 0.05 level (Tukey's method, adjusted for multiple comparisons).

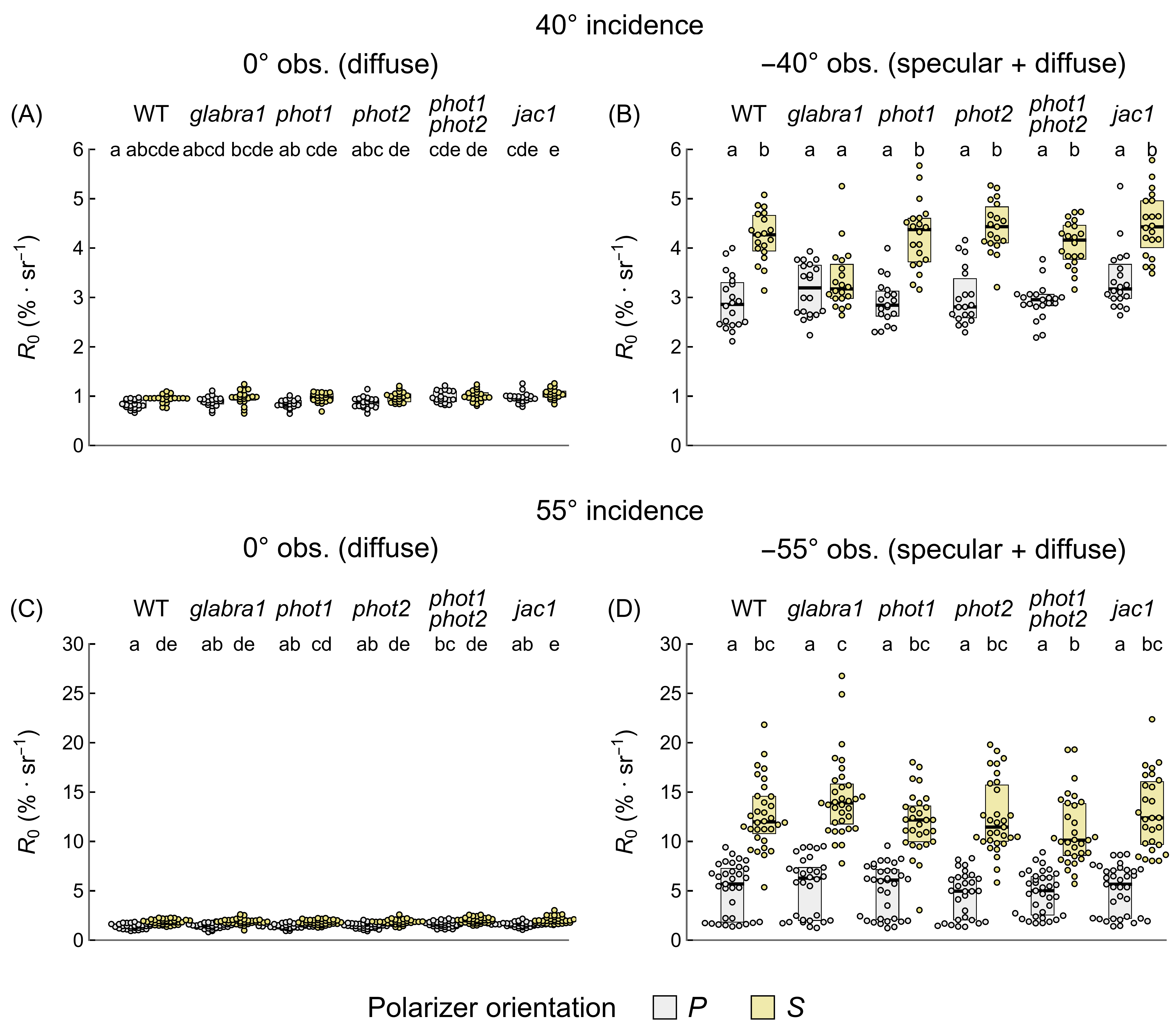

Fig. S8. Red light bidirectional reflectance of the adaxial side of leaves detached from dark-adapted *Arabidopsis* wild type (WT), *glabra1,* and chloroplast movement mutants (*phot1*, *phot2*, *phot1phot2*, *jac1*). The angle of incidence of unpolarized measuring light was either 40° (A, B) or 55° (C, D), while the angle of observation was set to 0° (diffuse reflectance recorded) (A, C) or either -40° (B) or -55° (D) (combined specular and diffuse reflectance recorded). The transmission axis of the detector polarizer was either parallel (*P,* grey boxes) or perpendicular (*S*, yellow boxes) to the plane of incidence. Each dot represents a value obtained for one leaf, with a single leaf harvested from each plant. Boxes show interquartile ranges, while the horizontal lines mark medians. Groups that do not share any letter differ at the 0.05 level in means of transformed values (Tukey's method, adjusted for multiple comparisons).

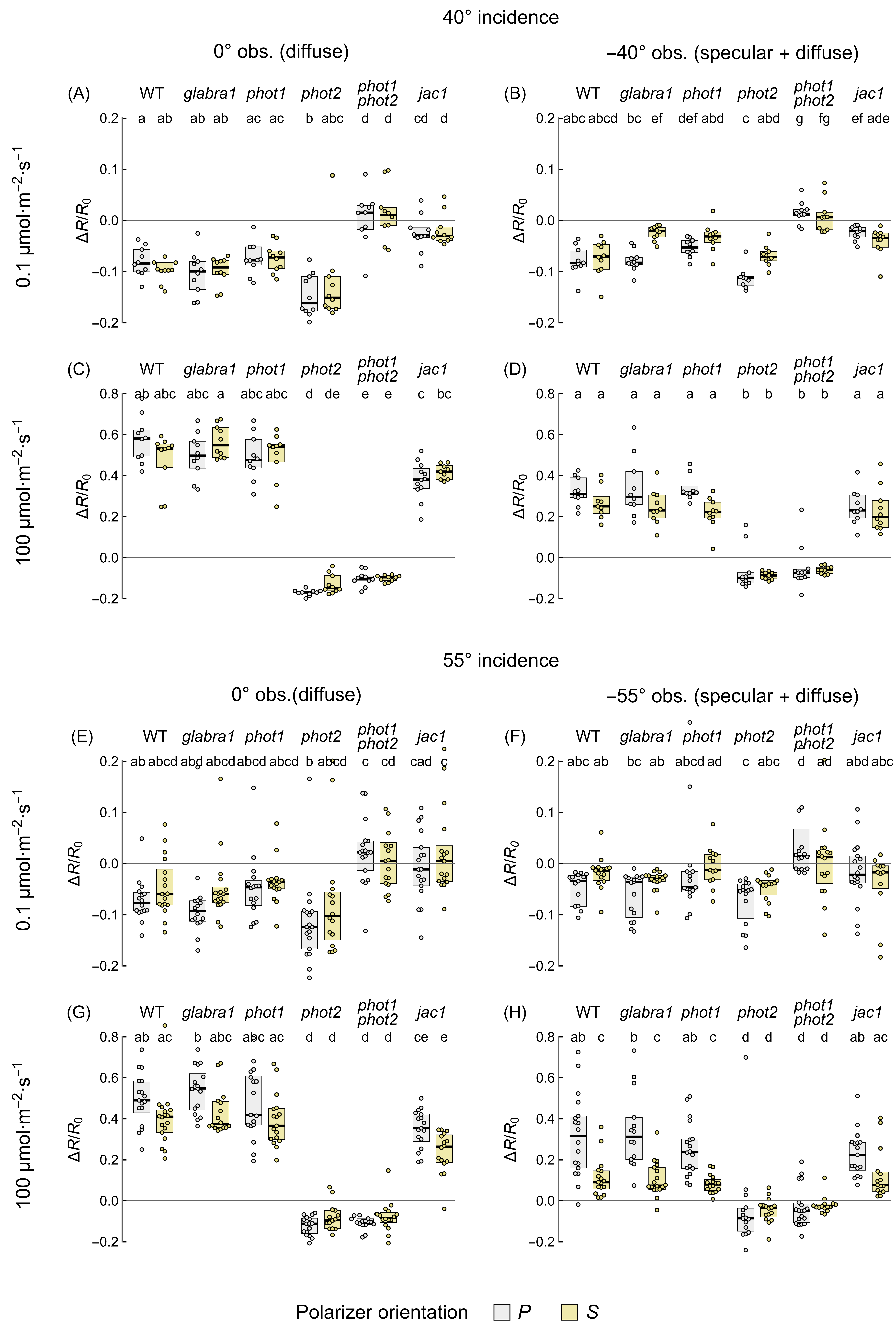

Fig. S9. The sensitivity of red-light leaf bidirectional reflectance to the chloroplast positioning, calculated as the ratio *ΔR/R_0_* between reflectance change *ΔR* induced with blue light of 0.1 (A, B, E, F) or 100 µmol·m^-2^·s^-1^ (C, D, G, H) and the initial reflectance *R_0_* of a dark-adapted leaf. The angle of incidence of actinic and measuring beams at the adaxial leaf surface was 40° (A - D) or 55° (E - H). The incident beams were not polarized. To measure diffuse reflectance (A, C, E, G), the detector was positioned at 0° to the leaf normal. To measure combined specular and diffuse reflection (B, D, F, H), the detector was positioned at -40° or -55°. For each detector position, two polarizer orientations were used: the transmission axis of the polarizer was either parallel to the plane of incidence (*P*, attenuates specular reflection, grey boxes) or perpendicular (*S*, yellow boxes). Each dot represents a value obtained for one leaf, with a single leaf harvested from each plant. Boxes show interquartile ranges, while the horizontal bars mark medians. Boxes that do not share any letter represent groups for which the means of transformed values differ at the 0.05 level (Tukey’s method, adjusted for multiple comparisons).

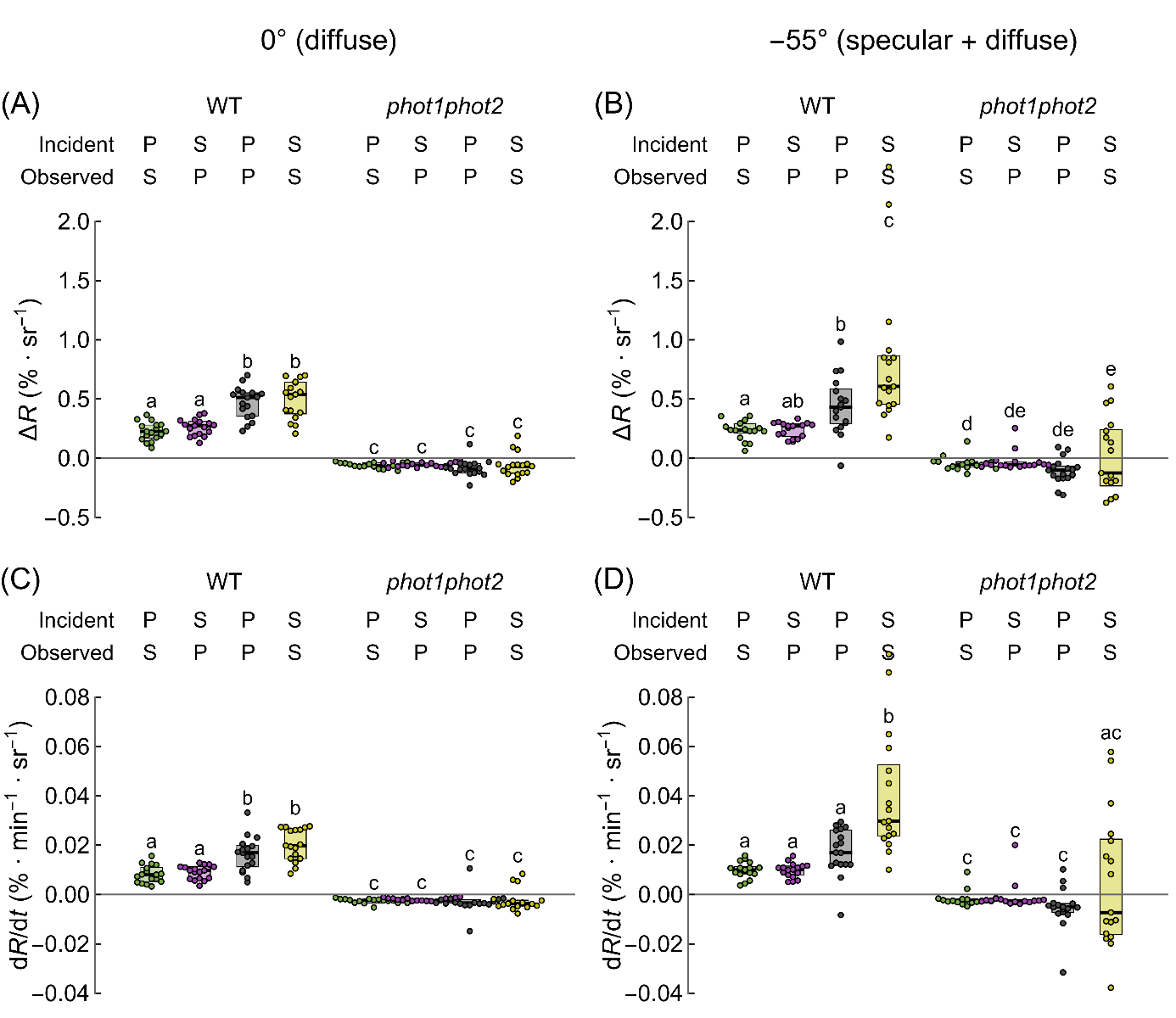

Fig. S10. Amplitudes *ΔR* (A, B) and rates d*R/*d*t* (C, D) of changes in the polarized red-light bidirectional reflectance induced by 1 h irradiation with high blue light (100 µmol·m^-2^·s^-1^) of the adaxial surface of leaves of wild-type (WT) and *phot1phot2* *Arabidopsis* plants. The blue actinic and red measuring beams impinged the adaxial leaf surface at the angle of 55°. For measurements of diffuse reflectance (A, C), the detector was positioned at 0° to the leaf normal. For measurements of combined specular and diffuse reflectance (B, D), the detector was positioned at -55°. Four possible combinations of the orientation (*P/S*) of the polarizers in front of the measuring light source and the detector were examined. Each dot represents a value obtained for one leaf, with a single leaf harvested from each plant. Boxes show interquartile ranges, while the horizontal bars mark medians. Boxes that do not share any letter represent groups for which the means of transformed values differ at the 0.05 level (Tukey's method, adjusted for multiple comparisons).

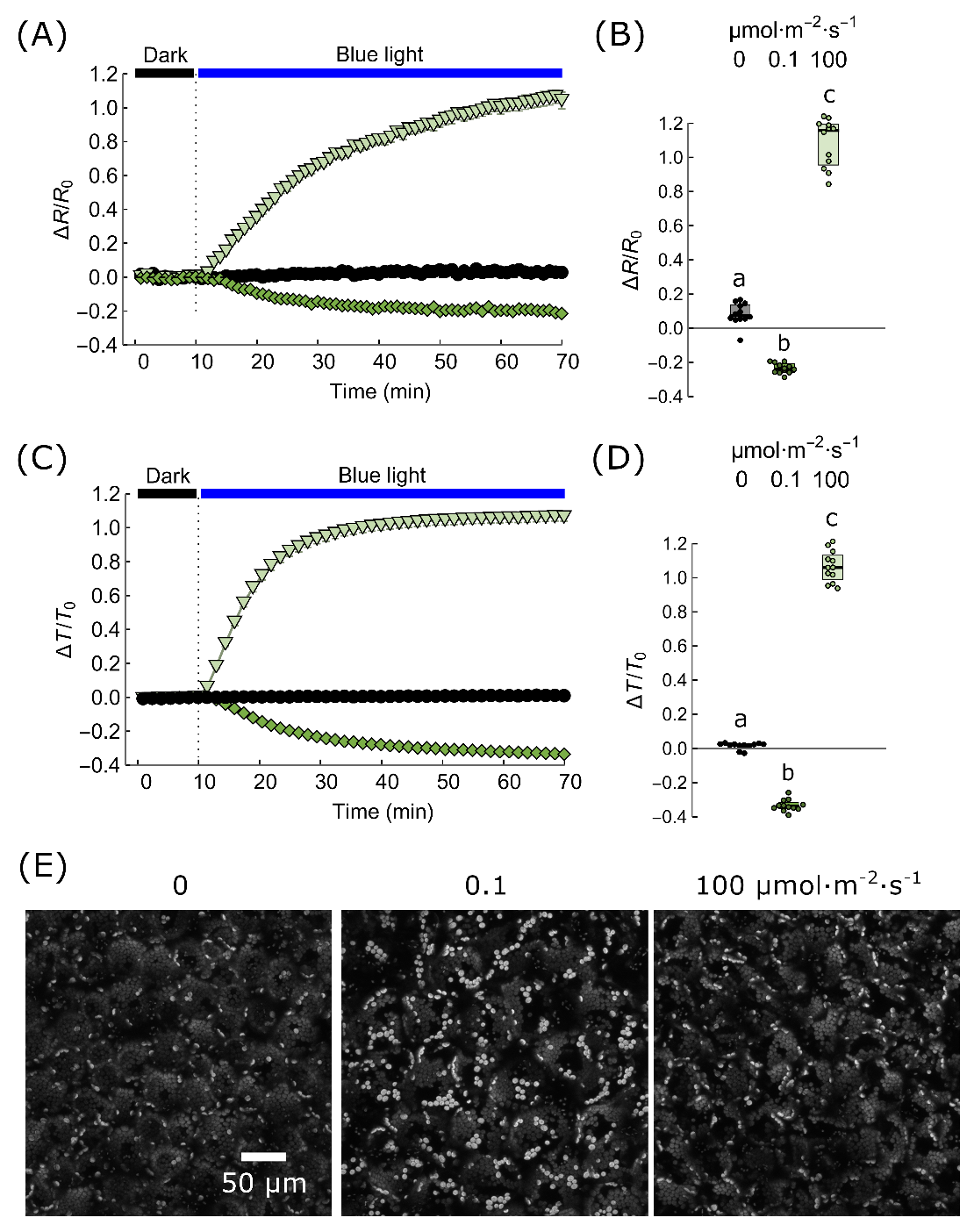

Fig. S11. Chloroplast movements examined with reflectometric (A, B), photometric (C, D), and microscopic (E) observations, in laboratory-grown *Arabidopsis thaliana*. Time courses of changes in red-light bidirectional reflectance (A) and hemispherical transmittance (C) induced by 1 h irradiation with low (0.1) or high blue light (100 µmol·m^-2^·s^-1^) or mock irradiation of the adaxial side of leaves. The angle of incidence of beams was 55°, while the angle of observation was 0° (diffuse reflectance recorded). The incident measuring light was polarized in the direction parallel to the plane of incidence (*P*). The polarizer in front of the detector (analyzer) transmitted light perpendicular to the incidence plane (*S*). Each dot represents a value obtained for one leaf harvested from a different plant. Boxes show interquartile ranges, while the horizontal bars mark medians. The means of transformed values from groups that do not share any letter differ at the 0.05 level (Tukey's method, adjusted for multiple comparisons). Confocal microscopy images (E) of leaf tissue after 1 h irradiation with low blue light (0.1 µmol·m^-2^·s^-1^) or high blue light (100 µmol·m^-2^·s^-1^) or mock irradiated.

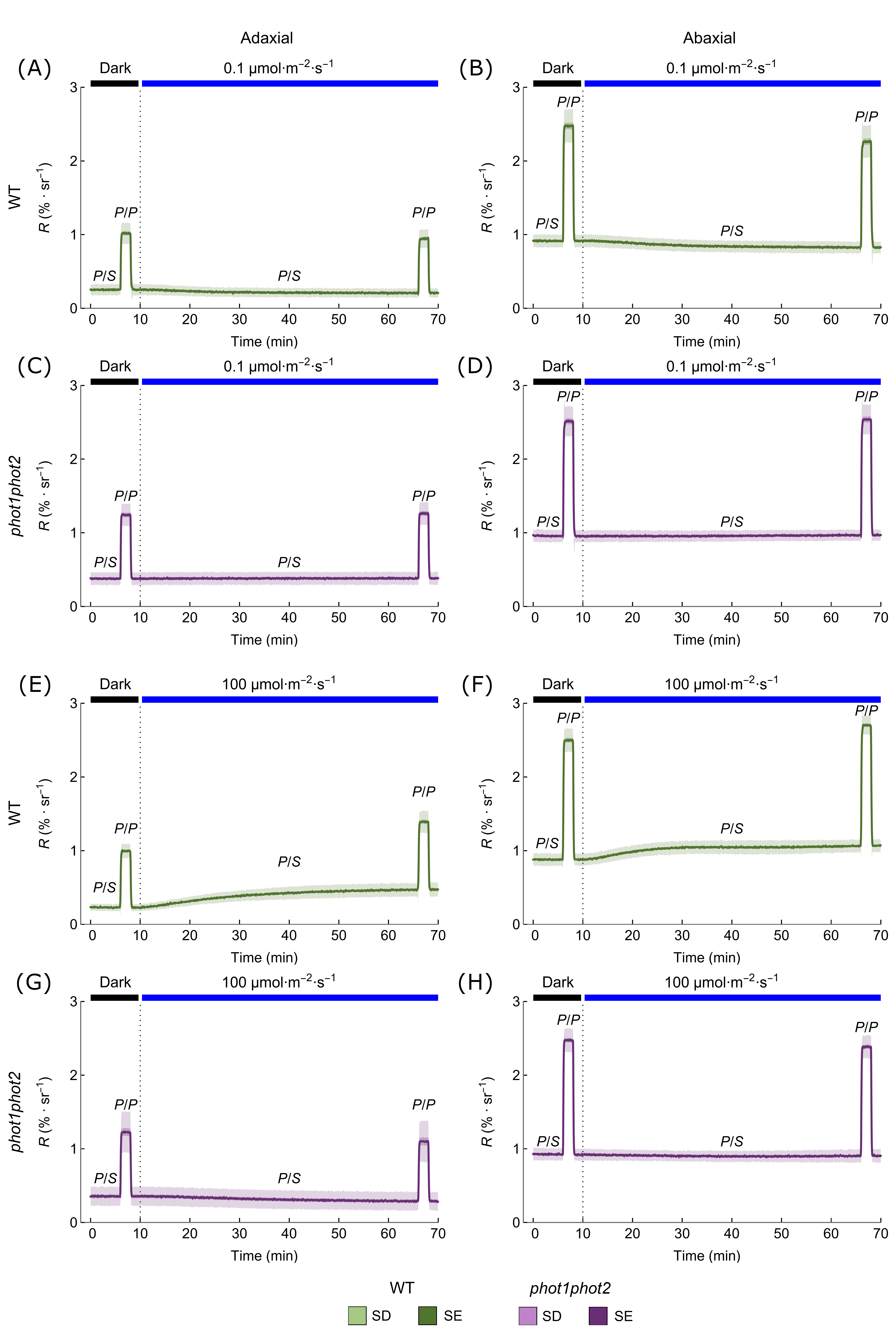

Fig. S12. Changes in the polarized red-light bidirectional reflectance induced by 1 h irradiation with blue light of 0.1 µmol·m^-2^·s^-1^ (A-D) or 100 µmol·m^-2^·s^-1^ (E-H)) of the abaxial and adaxial surface of leaves of wild type (WT) and *phot1phot2* *Arabidopsis* plants. The orientation of the polarizers in front of the measuring light source and the detector was changed during the experiment from the *P/S* position to *P/P*. The blue actinic and red measuring beams impinged the adaxial leaf surface at the angle of 55°. The detector was positioned at 0° to the leaf normal.

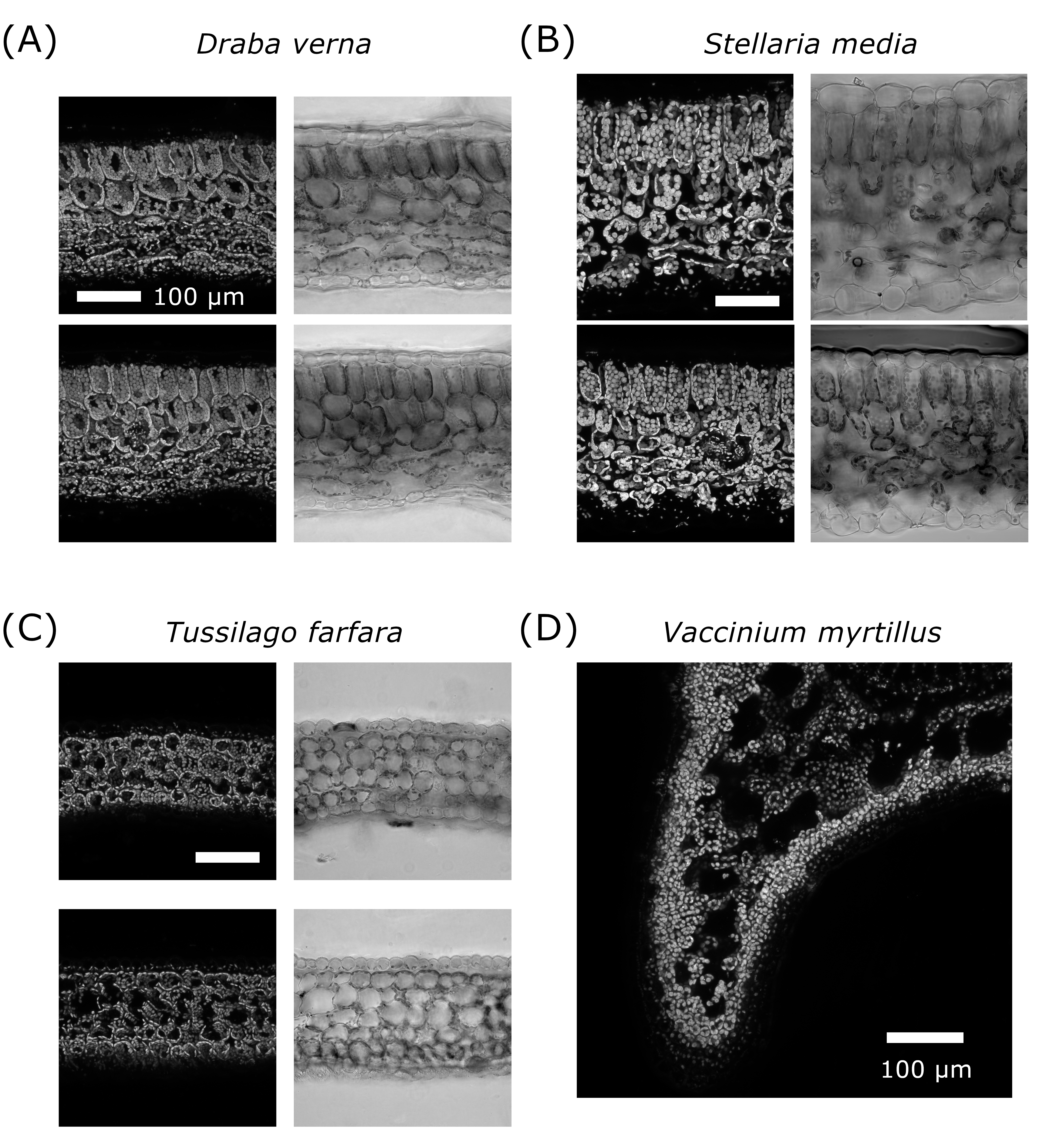

Fig. S13. Laser scanning confocal microscopy images of cross-sections *Draba verna* (A)*, Stellaria media* (B) leaves*, Tussilago farfara* (C) flowering shoot scales, and *Vaccinium myrtillus* (D) stem. Chlorophyll autofluorescence was recorded. Scale bars 100 µm.
